# Grandmother’s chemical legacy of pesticide exposure: bi-generational effects and acclimation in a model invertebrate

**DOI:** 10.1101/2020.11.25.397554

**Authors:** Rikke Poulsen, Henrik H. De Fine Licht, Martin Hansen, Nina Cedergreen

## Abstract

Man-made chemicals are a significant contributor to the ongoing deterioration of ecosystems. Currently, risk assessment of these chemicals is based on observations in a single generation of animals, despite potential adverse intergenerational effects. Here, we investigate the effect of the fungicide prochloraz across three generations of *Daphnia magna.* We studied both the effects of continuous exposure over all generations and the effects of first-generation (F0) exposure on two subsequent, non-exposed, generations. Effects at different levels of biological organization were monitored. Acclimation to prochloraz was found after continuous exposure. Following F0-exposure, non-exposed F1-offspring showed no significant effects. However, in the F2 animals, several parameters differed significantly from controls. A direct association between grandmaternal effects and toxic mode of action of prochloraz was found, showing that chemicals can be harmful not only to the exposed generation, but also to subsequent generations and that effects may even skip a generation.

## Introduction

The present age has been named the Anthropocene as humans are significantly impacting the Earths eco- and geosphere with loss of biodiversity, contamination of waterbodies, air pollution and deterioration of ecosystems as the result (*1*–*3*). Man-made chemicals are a significant contributor to these effects (*4*). Pharmaceuticals, fossil fuels, industrial chemicals, pesticides etc. are dispersed into the environment and these anthropogenic chemicals are almost omnipotent in today’s ecosystems as long-range transport and food-chain build up means that even pristine regions like the arctic or remote mountain ranges show measurable concentrations (*5*–*7*). Many of these chemicals are toxic and their presence causes harmful effects in humans and wildlife (*8*). For this reason, it is essential that studies of how animals interact with their environment in the twenty-first-century consider exposure to anthropogenic chemicals. Many countries are increasingly monitoring and regulating the dispersion of pollutants such as pesticides, biocides, and waste water components such as pharmaceuticals (*9*, *10*). Progressing from the initial focus on acute environmental effects resulting from lethal dosages, the focus of the last decades has increasingly been on long-term effects such as endocrine disruption and effects on reproduction (*8*, *11*). However, increasing scientific evidence of multigenerational effects suggests that the time perspective of the effect of chemical exposure has to be extended even further (*12*, *13*).

It is already documented that environmental cues in early development, other than man-made chemicals, can prepare the offspring for the expected environment in adulthood. For instance, when mothers of the water flea species *Daphnia cucullate* and *Daphnia galeata mendotae* are exposed to fish kairomones (chemical predator cues) they can produce daughters with a defensive “helmet” that prepares them to avoid predation. This expands the timeframe for when effects of chemical exposure can be observed. It also links chemical exposure to trade-offs in future generations, because if the experienced environment differs from the predicted, mal-acclimation will result along with a fitness consequence. For instance, the helmeted *Daphnia* have a smaller brood chamber, giving smaller clutches and lower reproductive rates (*14*). It is possible that similar biological events are happening with man-made chemicals and we might be missing an important point in how early organisms can be affected.

Currently, the only legal requirement in, for example, pesticide risk assessments (*16*) is to investigate one generation, sometimes with the inclusion of quantifying reproduction as number of offspring. The potential for multigenerational effects challenges the aptness of this status quo. If for instance effects of a chemical spill persist in several generations after the exposure has taken place, it will lead to a time gap and a misperception of correlations. Effectiveness of pollution remediation may be overestimated as intergenerational harmful effects continue after pollutants are removed from the ecosystem. Also, if prenatal life stages are sensitive to chemical exposure, it means that current risk assessments are missing a critical time point for evaluating effects of chemical exposure, as analysis of for example offspring fertility is not included in any current guidelines. To address this concern, this study considers an extended time perspective of toxicity. Applying *Daphnia magna* as the model organism, we examine the intergenerational effects of the well-characterized toxicant prochloraz over three generations.

*Daphnia magna* (Crustacea: Branchiopoda: Cladocera) is a much-favored model organism. They are ubiquitous in aquatic environments where they serve as an important link in the food chain (*17*). Furthermore, they reproduce by cyclic parthenogenesis, which provides an opportunity to separate genetic and environmental components of phenotypic plasticity (*18*, *19*). Prochloraz and other azole fungicides are approved for use in several EU countries (*20*), and have been found in the environment (*21*). Prochloraz is a well-characterized toxicant known to both block and induce cytochrome P450 monooxidase (CYP) enzymes (*22*), leading to endocrine disruption (*23*–*25*) and synergistic interaction with other toxicants (*26*–*28*). Following the adverse outcome pathway (AOP) framework, prochloraz toxicity in fish has been shown to result in the adverse outcome of reduced cumulative fecundity and decreased population trajectory (*29*). The key mechanisms behind these effects are the reduced concentration of 17β-estradiol and reduced vitellogenin synthesis linked to the molecular initiating event of CYP19, or aromatase, inhibition. CYP19 is a key enzyme in steroidogenesis and is responsible for the conversion of androgens to estrogens (*29*). In *Daphnia,* prochloraz is known to be a potent inhibitor of non-specific CYP enzyme activity *in vivo* (*30*), decrease reproduction, affect molting following long-term exposure (*31*), and lead to developmental abnormalities in offspring (*31*). However, wider generational effects of prochloraz, or any other chemical with the same mode of action, have not been investigated beyond F1 offspring abnormalities.

Here, we studied intergenerational effects across three generations of *Daphnia* resulting from prochloraz exposure, addressing both adverse effects as well as acclimation effects during and beyond exposure. Animals were followed through two exposure scenarios; 1) continuous exposure of all three generations (F0, F1 and F2) and 2) exposure of only the first generation (F0) (Figure 1A). We studied effects at different levels of biological organization combining genome-wide gene expression (using transcriptomics), whole organism metabolite levels (using metabolomics), and measurements of CYP enzyme activity as well as key phenotypic life-history effects, such as growth and reproduction.

**Figure 1.**
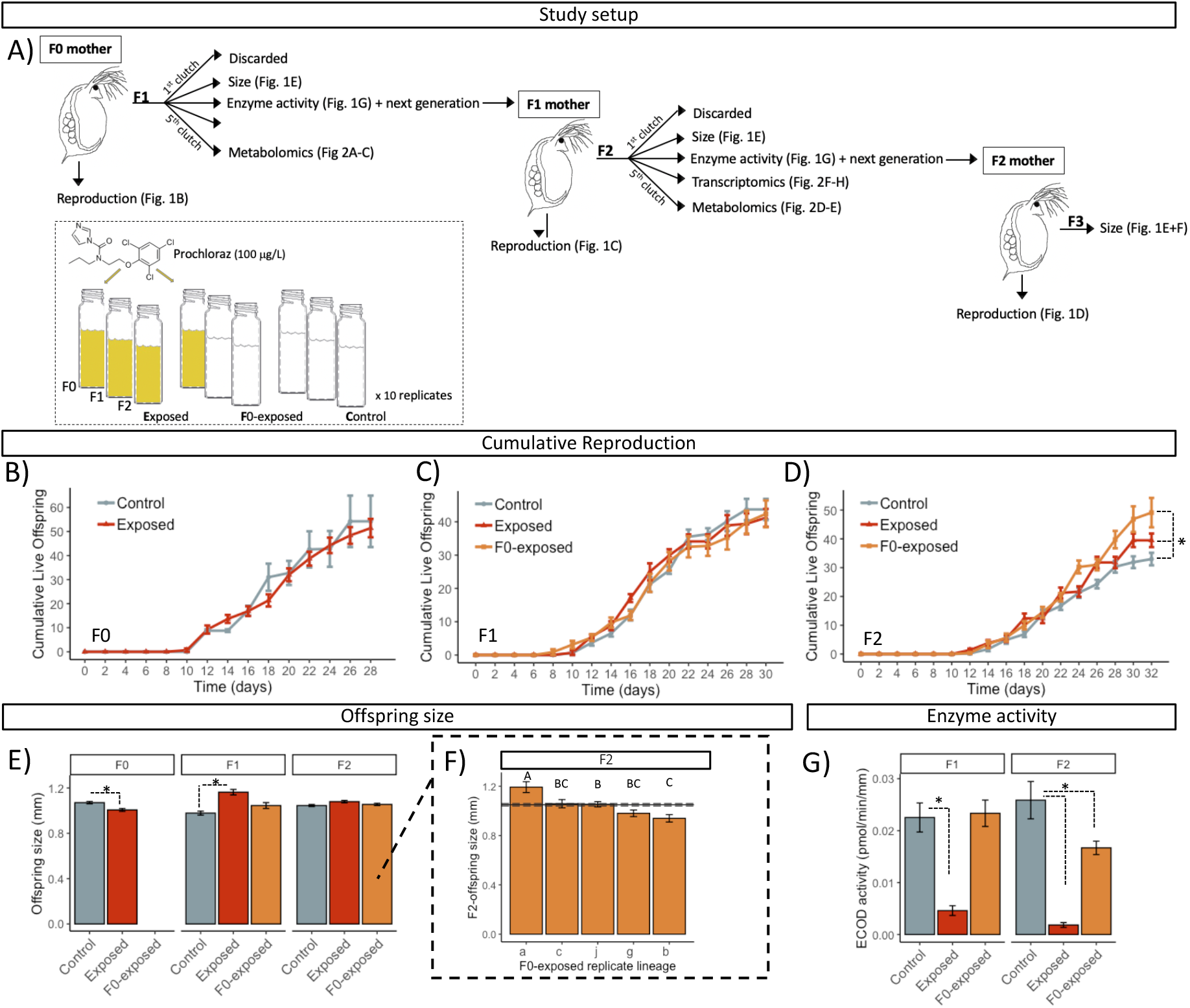
Intergenerational phenotypic and enzymatic effects of *D. magna* continuously exposed and only exposed in F0 generation to the fungicide prochloraz. **(A)** Experimental scheme. **(B-D)** Reproduction for the three treatment groups: control (blue), continuously exposed (red) and F0-exposed (orange) quantified as cumulative production of live offspring in generation F0 (B), F1 (C) and F2 (D). Error bars show standard error of the mean (SEM) (n=10 replicates), * indicate significant difference (Tukey′s post hoc *T-test p*<0.05). **(E)** Offspring length (mm) ± SEM (n=10 replicates for each treatment and N= 4-19 animals per replicate) in the three treatment groups and for the three generations F0-F2. * indicate significant difference between groups in the same generation (Tukey′s post hoc *T-test p*<0.05). **(F)** Lineage specific offspring length (mm) ± SEM in the F2 generation for the F0-exposed animals, plotted in the sequence that they occur in the PCA plot for gene expression (Figure 2F). The red line indicates the average length of the control animals and the dotted line SEM. Letters above the error bar show the statistical grouping (p-value < 0.05, N*=*30-61 animals per replicate). **(G)** Cytochrome P450 ECOD activity normalized to length of animals (pmol/min/mm) ± SEM (n=10 replicates for each treatment and N=10 animals per replicate) in control (blue), continuously exposed (red) and F0-exposed animals (orange) and for generations F1 and F2. *indicate significant difference (Tukey′s post hoc *T-test p*<0.05) from control group in the same generation.

## Results and discussion

### Acclimation after continuous prochloraz exposure

To investigate acclimation and compensatory responses over generations, we compared *Daphnia* lineages continuously exposed to prochloraz for three generations with control animals (Figure 1A, Treatment “Exposed” vs “Control”).

#### Effects on reproduction during continuous exposure

In the first generation (F0), exposure to 100 μg/L prochloraz significantly decreased offspring length when compared to controls (Tukey′s post hoc *T-test p*=0.002) (Figure 1E), while no difference was found in offspring amount (Figure 1B) (Tukey′s post hoc *T-test p*=0.7). This is consistent with our pilot study, where a 100 μg/L exposure corresponded to <*EC_1_* for cumulative offspring but >*EC_99_* for length (Figure S1A-C, Table S1). It is therefore evident that without pre-exposure the fungicide decreases length of offspring.

With continued exposure into generation F1 and F2, the average length of offspring was first significantly larger in F1 (Tukey′s post hoc *T-test p*<0.0001) but then similar to controls in F2 (Tukey′s post hoc *T-test p*=0.1) (Figure 1E). The size of offspring is important as larger offspring have better energy reserves to withstand environmental stress (*32*). The increase in size in F1 is therefore a likely compensatory response to continuous exposure to prochloraz, although larger size also comes with a higher risk of predation (*33*). Such changes in life history traits have usually been found to correlate with a change in per offspring investment (i.e. fewer but larger, or more but smaller offspring) (*34*). However, we observed a comparable number of offspring across treatments in F1 (Figure 1C). Such a lack of trade-off in *D. magna* reproduction was also observed for another azole fungicide, epoxyconazole, where both increased length and increased amount of offspring were observed after low-dose exposure (*35*). In addition, increased offspring production without effects on length was observed in response to two serotonin re-uptake inhibitors (*36*). The latter was later correlated with an increased aerobic catabolism and a higher sensitivity to anoxic conditions (*36*), showing trade-offs on other fitness traits than offspring number and size.

While cumulative reproduction remained unaffected in F1 (Figure 1C), it was significantly higher than controls in F2 (Figure 1D) (Tukey′s post hoc *T-test p*=0.7 and p=0.01, respectively). Within generation F2, these continuously exposed animals, furthermore, had their first clutch earlier compared to controls (Tukey′s post hoc *T-test p*=0.003) (Figure 1D). This can explain the higher cumulative number of offspring. At concentrations 5 times higher than those tested here, prochloraz has been observed to delay development leading to delayed reproduction (*31*). On the contrary, earlier onset of reproductive maturity was observed in response to compounds such as fish kairomones (*37*) and fluvoxamine (*38*). It should also be noted that some variation between generations in terms of cumulative reproduction was observed in controls (see supplementary material). Furthermore, there was significant offspring mortality in F0 of the exposed animals when compared to control lineages of the same generation (Figure S5) (Tukey′s post hoc *T-test p*<0.001).

#### Effects on overall CYP-enzyme activity during continuous exposure

*In vivo* 7-ethoxycoumarin-O-dealkylation (ECOD) activity was used as a measurement of cytochrome P450 monooxidase (CYP) biotransformation capacity. The assay is based on a broad-spectrum substrate and measures overall CYP activity (*30*). The ECOD activity was significantly reduced for the continuously exposed animals in both F1 and F2 (Tukey′s post hoc *T-test, p*-value < 0.001) (Figure 1G). There, furthermore, were no signs of acclimation as ECOD activity did not differ between the two generations.

Prochloraz has earlier proved a potent inhibitor of *Daphnia magna in vivo* ECOD activity (*30*), so a decreased activity was expected. CYPs have key functions such as detoxification of xenobiotics (*40*) and hormone regulation of molting (*41*).The fungicide is known to interact with CYPs (*22*) through coordination of its azole moiety lone pair to the heme iron that is present in the catalytic site of the enzyme (*39*). Hence, CYP-dependent pathways are the most likely targets for toxic mode of action of prochloraz in non-target organisms.

#### Effects on the*Daphnia* metabolome during continuous exposure

We used untargeted metabolome analysis to monitor changes of internal metabolites resulting from prochloraz exposure. In the untargeted metabolome analysis, a total of 2525 metabolites were detected that could be compared between treatments and generations (Figure 2B-D). The highest amount of differentiating metabolite concentrations was found in the continuously exposed F1-animals, which showed higher concentration of 421 compounds and less of 170 compounds compared to controls (Figure 2B) (FDR-adjusted p < 0.05). With continued exposure into generation F2, the number of differentiating concentrations decreased to a total of only 86 compounds (Figure 2D) (30 up- and 56 down-concentrated, FDR-adjusted p < 0.05). This corroborates the physiological- and life-history compensation and acclimation observed in life history traits and in CYP activity. Annotation was possible for 73 of the up- and down-concentrated compounds. 44 of these were endogenous metabolites that could be assigned to metabolic pathways (Table S4 and S5). In generation F1 a general up-concentration of compounds in metabolic pathways involved in biosynthesis of amino acids, carbohydrate metabolism and lipid metabolism (Table S4 and S5) was observed. This is consistent with the observed larger reproductive output observed in the continuously exposed F1 mothers (Figure 1E). Several compounds of differentiating concentration could also be annotated as tentative candidates belonging to metabolic pathways specifically connected to the interaction between prochloraz and heme-containing enzymes (*42*). Among them were estriol, which is a metabolite of 17β-estradiol, the enzymatic product of aromatase. Estriol was found slightly up-concentrated (logFC = 1.3, FDR-adjusted p < 0.05) in continuously exposed F1 animals, indicating a compensatory response in the hormone system. Also, an important pathway in the invertebrate lipid metabolism, α-linoleic acid metabolism (*43*), was found to be affected in this treatment group. α-linoleic acid as well as the two CYP-substrates: (9Z,11E,15Z)-(13S)-Hydroperoxyoctadeca-9,11,15-trienoate and (10E,12Z,15Z)-(9S)-9-Hydroperoxyoctadeca-10,12,15-trienoic acid, were found in higher concentrations. A CYP-catalyzed reaction can convert the latter metabolite to (9S)-(10E,12Z,15Z)-9,10-Epoxyoctadecatri-10,12,15-enoic acid and this compound was found in lower concentration in exposed animals. This increased substrate concentration and decreased product concentration is a clear indication of CYP-enzyme inhibition.

**Figure 2.**
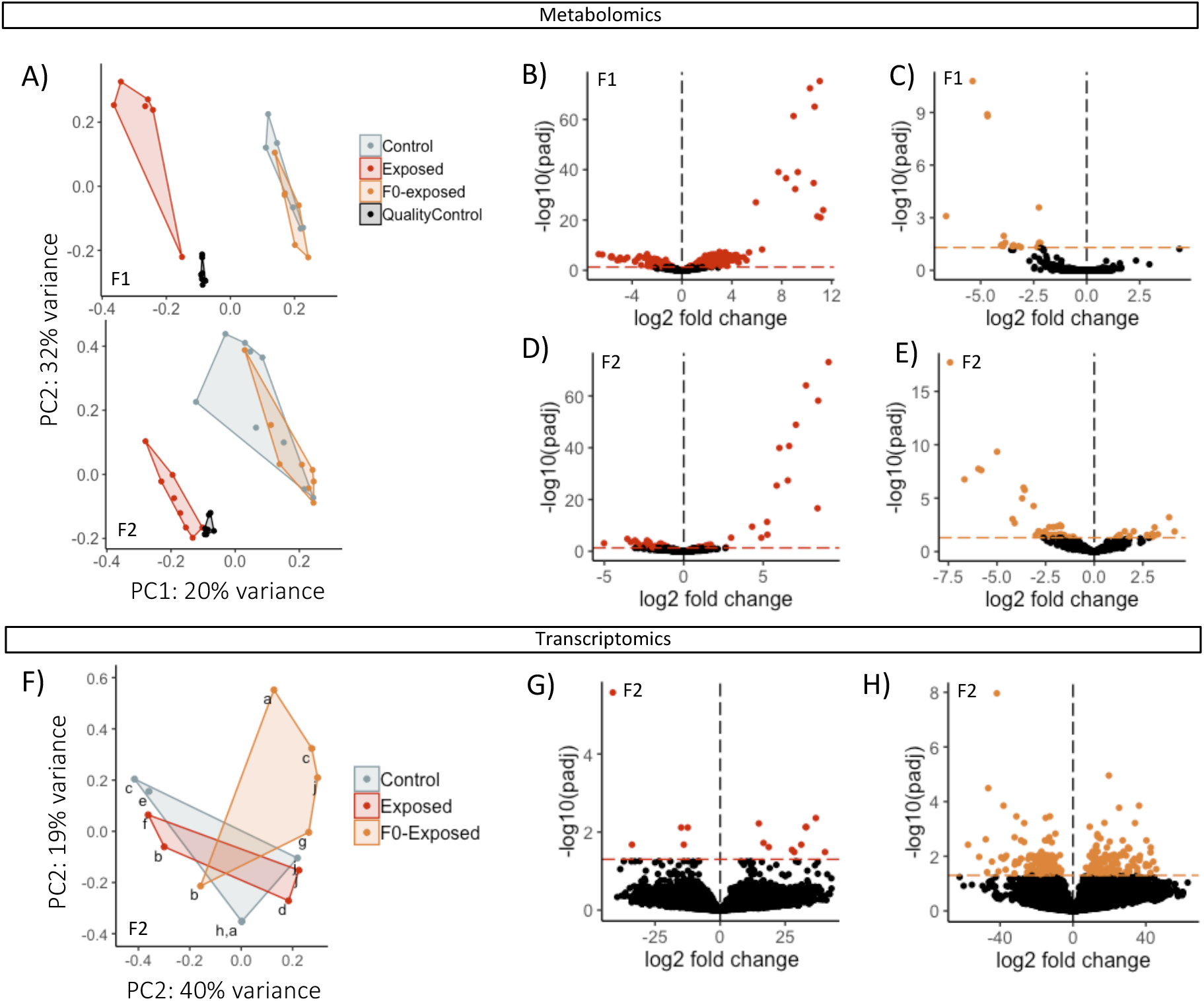
Intergenerational changes in genome-wide gene expression and metabolite levels in *D. magna* following continued and F0-exposure to procholraz. **(A)** Principal component analysis (PCA) for global metabolome analysis in generation F1(upper) and F2 (lower) for controls (blue), continuously exposed (red) and F0-exposed animals (orange). Dots indicate individual replicates, each consisting of metabolite extracts from 2-15 5-day old juvenile *D. magna*. Black dots indicate 7 replicates of the composite quality control sample. **(B-E)** Volcano plots showing down- and upconcentrated metabolites in 5-day old *D. magna* in continuously exposed F1 (B) and F2 (D) animals and in F0-exposed F1 (C) and F2 (E) animals. The x-axis represents positive and negative log2-fold changes, respectively. Metabolites highlighted in red/orange differ significantyly in their concentration when compared to control animals (p.adj<0.05). **(F)** PCA for gene expression in generation F2 for controls (blue), continuously exposed (red) and F0-exposed (orange) animals. Dots with letters indicate individual replicates, each consisting of RNA-extract from 5-10 5-day old juvenile *D. magna* **(F-G)** Volcano plot of F2-RNAseq data showing up- and down-regulated genes in 5-day old *D. magna* in continuously exposed F2 (F) and F0-exposed F2 (G) animals. The x-axis represents positive and negative fold changes, respectively) Genes highlighted in red/orange are significantly differentially transcribed compared to control animals (p.adj<0.05).

Effects were also observed on other compound concentrations that are controlled by heme-containing enzymes. This included compounds in the cutin, suberine and wax biosynthesis pathway, which for instance are involved in cuticle formation and metabolism of terpenoids involved in synthesis of insect hormones (Table S5). The affected metabolites add evidence to the molecular initiating event (MIE) of prochloraz toxicity being coordination of the lone pair of the azole moiety to the heme iron present in catalytic sites of enzymes. However, it also underlines that this can happen for several enzymes involved in many different pathways and not just for the aromatase with subsequent effects on the steroidogenesis.

Metabolites that were affected in F1 of continuously exposed animals were rarely affected in F2 or the effect was of opposite direction (Table S4 and S5). For instance, there was more steroid tetrahydrocorticosterone in F1 (log2 FC=2.2, FDR-adjusted p < 0.05) but less in F2 (log2 FC=-1.8, FDR-adjusted p < 0.05). This is a further substantiation of the intergenerational compensatory response and acclimation in *Daphnia* continuously exposed to prochloraz.

Prochloraz and its metabolites were not separated from the endogenous metabolites in the data processing. Thirteen prochloraz metabolites were putatively annotated (Table S6) and transformations, such as partial loss of the imidazole ring with subsequent aldehyde formation corresponded well with previous studies of prochloraz metabolism in the freshwater invertebrate *Gammarus pulex* (*40*)

#### Effects on the transcriptome in generation F2 after continuous exposure

Analysis of the transcriptome in F2 showed considerable inter-treatment variance (Figure 2F), and notably the overall gene expression in continuously exposed lineages was very similar to control lineages (Figure 2G). Three generations of continuous exposure to prochloraz gave only 10 up-regulated and 5 down-regulated genes (FDR-adjusted p < 0.05) (Figure 2G). Among these was, however, an upregulation of aromatase (GeneID: DMV1G132140T0, log2 fold change = 33, padj=0.007), which confirms that this enzyme plays a central role in prochloraz toxicity, also after continuous exposure in three generations. Taken together, all our data is consistent with intergenerational compensatory response and acclimation as a result of constant prochloraz in *Daphnia*. Such observed acclimating processes are likely to result in ecologically relevant trade-offs and have consequences for the ability of the animals to handle other challenges such as predation, other xenobiotics and anoxic conditions.

### Inter-generational effect of F0-exposure

After establishing that continuous prochloraz exposure has substantial physiological and metabolomic effects on Daphnia, we investigated whether toxic effects persist beyond the exposed individual and if ancestoral exposure influences subsequent un-exposed generations. This was done by comparing generation F1 and F2 of F0-exposed lineages with control animals (Figure 1A, Treatment “F0-exposed” vs “Control”).

#### Effects on reproduction after F0-exposure

Following the exposure in F0, which resulted in a decreased offspring length (Figure 1E), the non-exposed animals in the subsequent F1- and F2 generations produced offspring of similar size to control animals. However, in F2 the animals had significantly more offspring compared to both controls and to continuously exposed animals (Tukey′s post hoc *T-test p*<0.0001 and p=0.004, respectively) (Figure 1D). Equal to the increased offspring size observed in F1 of continuous exposed lineages, this increase in number of offspring supports a compensatory response in F2 to prochloraz exposure in F0.

#### Effects on overall CYP-enzyme activity after F0-exposure

The F0 exposed animals did not display any significant decrease in ECOD activity in the F1 generation, indicating full recovery from their mother’s exposure. However, when activity was measured in the F2 generation, it had decreased to a value intermediate between the control and continuously exposure animals (Tukey′s post hoc *T-test p* = 0.01 compared to controls of same generation) (Figure 1G). These results suggest that grandmaternal (F0) exposure leads to more pronounced effects on enzyme activity in granddaughters (F2) than in daughters (F1).

#### Effects on the Daphnia metabolome after F0-exposure

Pairwise comparison by multivariate statistical analyses showed differences from controls in both generation F1 and F2 (FDR-adjusted p < 0.05). Opposite to the continuously exposed animals, however, the largest differences were observed in F2. In F1, 21 metabolites were at lower concentrations when compared to controls and none were found up-concentrated (Figure 2C). In F2, 41 and 16 compounds were at lower and higher concentrations, respectively, when compared to controls (Figure 2E). This supports the observations made for both ECOD activity and cumulative reproduction, which showed a larger effect in F2 compared to F1. The metabolites that changed concentrations in F0-exposed animals included, for example, tetrahydrocortisol in generation F2 (log2 FC=1.8, FDR-adjusted p < 0.05) (Table S5). This metabolite is involved in biosynthesis of steroid hormones, which actually was equally affected in F1 of the continuously exposed animals (log2 FC=2.8, FDR-adjusted p < 0.05). Juvenile hormone III acid, a metabolite involved in insect hormone biosynthesis and a CYP product, was also affected in F2 of the F0-exposed, but not in F1. In the continuously exposed animals, this metabolite was affected in both generations. This supports the observation of CYP-activity being affected in grand-maternally exposed (F2), but not in maternally exposed (F1) animals.

#### Effects on the transcriptome in generation F2 after F0-exposure

The F2 gene transcription in grand-maternally exposed animals showed a much larger divergence from the controls than the continuously exposed animals (Figure 2F+H). In total, 170 genes were less expressed and 127 were more highly expressed (Data S1). Enrichment analysis of gene ontology (GO) terms among the 227 annotated differentially expressed genes using the hypergeometric distribution revealed four GO-terms that were significantly enriched (Bonferroni-adjusted P<0.05) in F0-exposed animals relative to controls in the F2 generation (Table 1, Table S7). All enriched GO-terms were related to oxidation-reduction processes and most also to iron-ion containing enzymes. Among the upregulated monooxygenases were four putative CYP-genes coding for CYP4aa1, 4d20 and 6a18; all part of pathways for metabolizing xenobiotics. Furthermore, the aromatase gene was significantly upregulated (Table 1). Significantly depleted GO-terms were related to the spliceosomal complex, RNA binding and mitochondrial transcription (Table S8). Thus, the functional characteristics of upregulated genes are directly related to the chemical characteristics of prochloraz and the well-documented interaction between the azole moiety and heme-iron (*39*). Hence, we show that the azole moiety and heme-iron is the molecular initiating event in an adverse toxic outcome of prochloraz. In addition, our data suggest that the same mode of action is affected across generations and in generations beyond the exposed generation in the case of a singular exposure event.

**Table 1.**
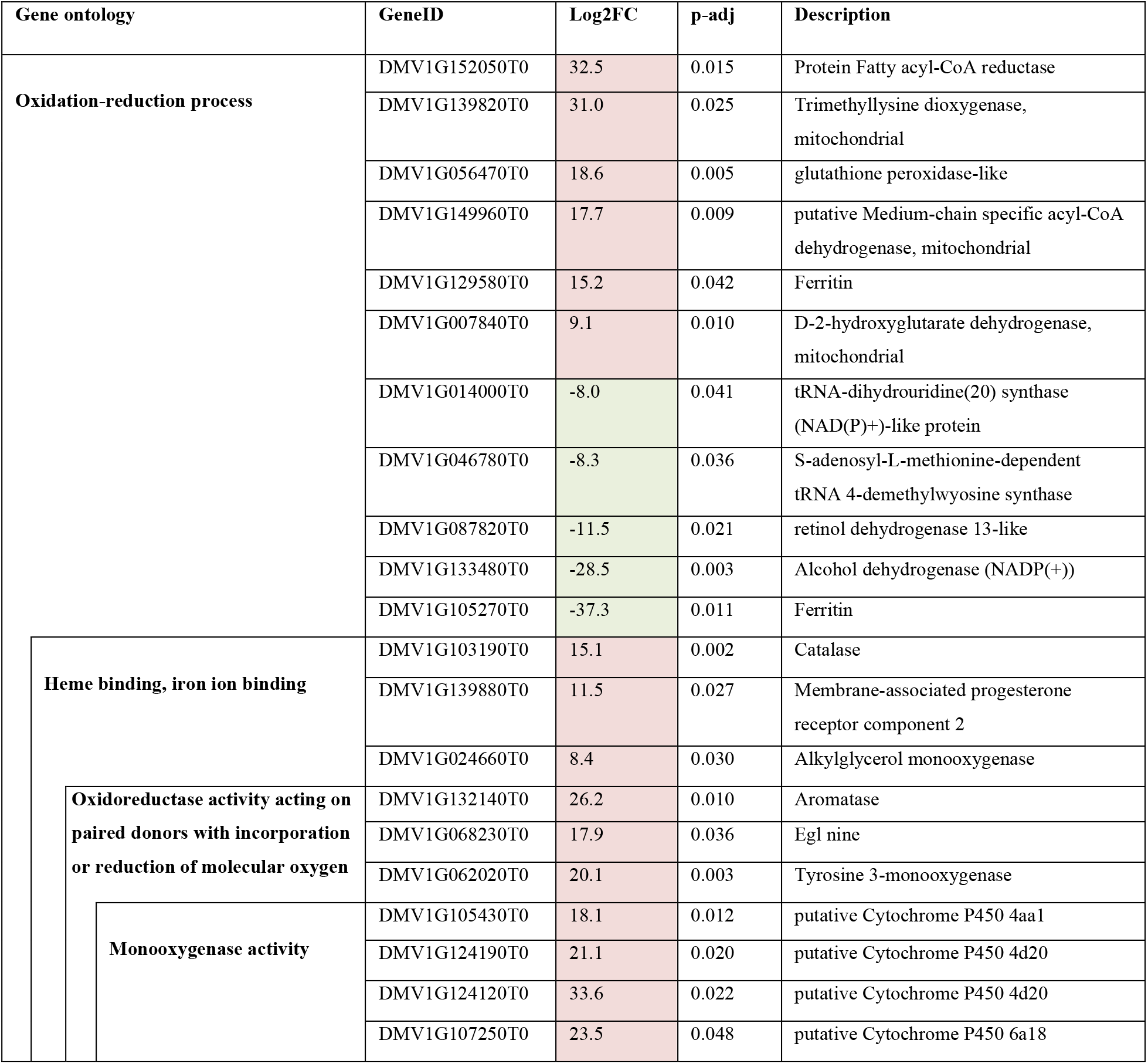
Significantly enriched gene ontology (GO) terms in F0-exposed animals compared to controls using the hypergeometric distribution (Bonferroni-adjusted P<0.05). The enriched GO-term, GeneID, log2 fold change and adjusted p-value for the statistical comparison, and finally the differentially expressed genes within this GO-category.

Furthermore, the effect on aromatase links the effect in *Daphnia* to the currently accepted AOP for prochloraz in vertebrates. It is, thus, evident that several CYP-dependent biochemical pathways may be affected by the fungicide, similar to the metabolome analysis in generation F1 of the continuously exposed animals.

### Are we overlooking the importance of grandmaternal exposure?

A grandmaternal effect of prochloraz, which we here define as significant differences between controls and F0-exposed animals in F2, but with no difference observed in F1, was found for the cumulative number of offspring (Figure 1D), cytochrome P450 activity (Figure1G), and up-concentration of metabolites (Figure 2E). It was further underlined by differential expression of several key genes in F2 (Figure 2H), which could be directly related to the mode of action of prochloraz (Table 1). This demonstrates that effects of prochloraz exposure in *D. magna* can last beyond the exposed generation and that there can be a time gap between the exposure and the emergence of effects.

Furthermore, it begs the questions of whether primordial germ cells are an important life stage to consider when evaluating effects of chemical exposure and assessing their risks. During the continuous exposure of the F0-generation, the F2 animals were present as primordial germ cells (PGCs) inside the developing F1s. Effects in F2 could, therefore, arise both because of a direct exposure of the PGCs or indirectly because of an intergenerational transfer of biological information from the developing embryo (F1) to the PGCs (F2). The PGC stage is a sensitive time point, especially for establishment of epigenetic marks that later are essential for the dynamic regulation of gene expression underlying the cellular plastic response to environmental- and developmental cues (*44*). Effect of chemicals on the epigenetic landscape of PGCs has for instance been observed in male mice where F0-exposure to the anti-androgenic fungicide vinclozolin changed the expression of ncRNAs in PGCs leading to histopathological effects in testes in F1-F3 (*45*). Epigenetic effects were not evaluated in the present study but several genes coding for epigenetically relevant genes were among the differentially expressed genes in F2 when comparing F0-exposed animals to controls (Table S9).

### Individual plasticity

The principal component analysis revealed an interesting pattern in the F0-exposed samples as the individual lineages were gradually more divergent form the controls (Figure 1F). Serially passaged line “F0-exposed_a” represented the largest distance while “F0-exposed_b” was most similar. This suggests that their gene expression represents different degrees of divergence from the baseline (control), or in other words inter-lineage plasticity within the F0-exposed treatment group. Interestingly, this pattern in gene-expression variation correlates with lineage-specific off-spring size within the F2 generation of the F0-exposed lineages. This is visualized in Figure 1F, where the length of the F2 offspring has been plotted in the sequence observed in the PCA plot. Since length was measured on several offspring, the variance within each replica is known and we were able to do statistical comparisons between the different lineages. The length is decreasing in the order observed in the PCA plot with statistically significant differences (p-value < 0.05). The continuously exposed lineages and the control lineages, which also appear in the PCA plot, did not differ significantly in length of F2 offspring. Since we assume that the clonal animals of the same lineage are so similar that they can be used as proxy for each other, it follows that transcription measured in these 10 5-day old animals will reflect the gene transcription in the individual selected to be the F2 mother of that lineage. The correlation between the transcription in F2 and the length of their 2-day-old offspring (F3) (Figure 1A), therefore, reflects the correlation between gene transcription in the mother and the length of her offspring. The pattern shows that the more similar gene expression is to the control animals, the smaller the offspring gets. This demonstrates that a gradual plastic adjustment is taking place and that there seems to be a metabolic cost in returning to control levels after prochloraz exposure, i.e. a trade-off between offspring size and the ability to cope with grandmaternal prochloraz exposure. Finally, we see that the response is not synchronized between the replicate lineages. *Daphnia* are known to exhibit a high degree of plasticity to various environmental stimuli (*18*, *19*, *32*). Consequently, it is likely that different isolated lineages in experimental setups, such as the current one, will lead to plastic divergence.

### Toxicity is a process

As exemplified, the molecular and physiological effect of chemical exposure reaches to subsequent generations of the exposed individual; whether it is a lingering effect in granddaughters or acclimation to the constant exposure. This implies that current risk assessment procedures of for instance pesticides could be neglecting an important aspect of toxicity. It also adds to the concern that toxicity observed in the field might be uncoupled from the measured chemical exposure. Finally, it opens new interesting questions about *Daphnia* developmental biology. As stated by Tjalling Jager in his paper “Predicting environmental risk: A roadmap for the future”: *“Toxicity is not a value, it is a process”*(*46*). An important part of this process is timing and the dynamics here are still not well understood. Future work should focus on multigenerational effects of chemicals and sensitivity of prenatal life stages to elucidate their importance for hazard assessment of chemicals in order to further improve adequate risk assessment of environmental pollutants and protect ecosystems

## Materials and Methods

### Chemicals and reagents

Prochloraz (CAS; 67747-09-5, purity 98.6%) and d4-prochloraz (purity 98%) were obtained from Sigma-Aldrich, Germany. The acetonitrile (CAS; 75-05-8, LC-MS grade) used for prochloraz quantification was obtained from Rathburn (Walkerburn, Scotland), acetic acid (CAS; 64-19-7, LC-MS grade) from Sigma-Aldrich (Germany), 25:24:1 phenol:chloroform:isoamyl alcohol from Carl Roth GmbH & Co. KG (Germany) and isopropanol (CAS; 67-63-0, >96% purity) and chloroform (99-99.6%, stabilized with 0.6% ethanol) used for RNA extraction from VWR (Denmark). MilliQ water was purified on a PURELAB Chorus system (ELGA LabWater, UK). Solvents used for extraction and analysis of metabolites including water (CAS; 7732-18-5), water with 0.1% formic acid, water with 0.1% trifluoroacetic acid, acetonitrile (CAS; 75-05-8), methanol (CAS; 67-56-1) and 2-propanol (CAS; 67-63-0) were all of OptimaTM LC/MS Grade and obtained from Fisher Scientific (Ontario, Canada). Dichloromethane (CAS; 75-09-2) was obtained from Rathburn (Walkerburn, Scotland). The reagents 7-ethoxycoumarin (CAS: 31005-02-4, purity: 99.7 %) and RNAlater®was purchased from Sigma-Aldrich (Germany), TRIzolTM reagent from ThermoFisher Scientific, (USA) and diethyl pyrocarbonate (DEPC, CAS; 1609-47-8) from MP biomedicals (Germany).

### Culture conditions

The *D. magna* (Crustacea: Branchiopoda: Cladocera) clone was originally isolated from Langedammen, Birkerød, Denmark in 1978. Culturing was performed as described in Kretschmann et al., (2015). In brief the animals were cultured in M7 medium (*47*) and fed with the green algae, *Raphidocelis subcapitata*, (approx. 9.0 × 104 cells/(daphnid·d)) which had been cultured according to the ISO guideline no. 28692 (1989). At study initiation, offspring (≤24 h) from 30 4-week old mothers were separated, pooled together and randomly chosen individuals were assigned to 60 mL clear glass vials closed with a white plastic lid with an aeration hole stopped with a pipette filter tip. Culturing condition remained the same except for food amount, which was modified to age (Table S1).

### Study design

Figure 1A illustrates the experimental design. With the general setup based on the OECD guideline for *Daphnia magna* reproduction test (*48*), animals were followed for 3 generations. There were three treatment groups; one where all generations were exposed to prochloraz (Exposed), one where only the first generation was exposed (F0-exposed) and finally a non-exposed control group (Control). For each treatment in each generation there were 10 replicate glasses all containing one mother, resulting in 10 3-genertion lineages per treatment. In generation F0, the continuously exposed lineages (Figure 1A, Treatment “Exposed”) and the F0-exposed (Figure 1A, Treatment “F0-exposed”) experienced the same exposure scenario and the number of replicates here are therefore 20. This sample size was not determined based on statistical methods.

The endpoints assessed in the experiment ranged from physiological effects, with emphasis on growth and reproduction, to enzyme activity measurements, transcriptomics and metabolomics (Figure S1). Unfortunately blinding was not possible for the study. *D. magna* produce offspring in distinct clutches and the different offspring clutches were used in different analyses. Hence, we made the assumption that clonal animals of the same lineage were so similar that they could be used as proxy for each other. The first clutch was only counted and added to total number of offspring. The second clutch was used for length measurements, and subsequent clutches were either used to start the next generation or for endpoint measurements. In order to achieve sufficient biomass and make the setup comparable with the enzyme assay protocol, measurements of the molecular endpoints (i.e. ECOD, transcriptomics and metabolomics) were performed on 5-day old juveniles: 10 offspring from the same replica and line were reared to day 5 in a 250 mL blue cap bottle containing 200 mL of a test solution comparable to the one that the following generation was reared in (i.e. juveniles of the treatment group “F0-exposed” were transferred to pure M7 medium). They were fed ad libitum and treatment and medium was not changed during the 5 days.

### Exposure

The *D. magna* mothers were kept in 60 mL vials containing 50 mL exposure medium. The concentration of prochloraz was set to 100 μg/L based on a pilot study of the dose-response relationship (Figure S1) according to the OECD guideline for 21-day reproduction study in *D. magna*. The selected dose affected length of offspring but not the number of offspring and it strongly inhibited cytochrome P450 ECOD activity. Reported environmental concentrations of prochloraz are below 1 μg/L (*21*), which is a factor of 100 lower than the targeted exposure in the current study. The environmental relevance of the reported effects is therefore not the goal of this paper. The test solutions were mixed in blue cap flasks just prior to exposure renewal by adding 8 mL stock solution (6.25 mg/L prochloraz in MilliQ water), algae suspension (see supplementary information for feeding scheme), followed by M7 medium to a total volume of 500 mL. Test solutions were renewed every two days.

### Exposure verification by UPLC-MS/MS

In order to monitor actual exposure concentrations, water samples were collected from three random vials within each treatment group before change and from each of the new test solutions before distribution into test vials. Prochloraz concentrations were quantified using UPLC-MS^2^ with deuterated prochloraz (d4-prz) as internal standard.

#### Sample preparation

The M7 medium contains many different salts, which may influence the ionization of the MS by causing ion suppression or enhancement (*49*, *50*). A 20x dilution was therefore used as sample preparation to decrease these matrix effects below 10%. Initially samples were centrifuged at 14000 g for 5 minutes in order to precipitate larger particle. 50 μL of the cleared supernatant was combined with 950 μL mobile phase A (purified water with 0.01% acetic acid (v/v) and 5% acetonitrile) fortified with 105 ug/L internal standard (IS). In this way the final IS concentration in the vials was 10 ug/L.

#### Liquid chromatography

Chromatographic separation was performed on a C_18_ analytical column (Acquity UPLC BEH, 1.7 μm C_18_, 130 Å, 50 × 2.1 mm, Waters, USA) with a guard column (VanGuard, C_18_, 2.1 mm, Waters, USA) placed before the analytical column. The thermostated auto sampler was set to 10 °C and 5 μL of sample was injected on a heated column (40°C), using a mobile phase A composed of purified H_2_O with 5% acetonitrile and a mobile phase B containing 100% acetonitrile. With a flow rate of 0.450 mL/min the elution gradient was maintained at 10% B for the first 2 min, 10.0–95.0 % B from 2.0 to 2.2 min, held at 95.0 % B from 2.2 to 5.2 min, before re-equilibrating the column by the decrease 95%-10% B from 5.2-5.3 and held at 10% B for 1.7 minutes. The total method lasted 7 min yielding retention times (Rt, min) of 2.82.

#### Mass spectrometry

Detection was performed with a WatersXevo TQD triple quadrupole mass spectrometer with electrospray ionization (Waters, Milford, USA). Based on the method by EU reference laboratory for pesticides (*51*) and taking advantage of the isotopic pattern for chlorine, the ion transitions (m/z values) were 376>308 and 378>310 for prochloraz, and 380>312 and 382>314 for d4-prochloraz. Parameter settings for the mass spectrometer were optimized by injecting a 50 μg/L standard of prochloraz and d4-prochloraz directly in the MS, including LC flow with eluent composition as at the point of elution. Optimal cone voltage and collision energy were found using the software Intellistart (Waters, Milford, USA). The ion source was run in positive mode and kept at 150 °C with a capillary voltage of 3.5 kV and cone voltage of 22V. A desolvation gas flow of 600 L/hr at 500 °C and a cone gas flow of 20 L/hr were applied. To protect the MS from remaining salts and other early eluting impurities, a method event sequence was included, directing the LC flow to waste for the first 2.2 min, and again from 3.2 min onwards. Collection- and data treatment was performed with Waters MassLynx ^TM^ software (v4.1, Waters, Millford, USA)

#### Method validation

Matrix effects were evaluated by adding different amounts of M7 medium to neat standards of prochloraz. It was found that a 20 times dilution was needed to decrease the suppressing matrix effect to 10% and consequently samples were diluted 20 times in mobilephase A as sample preparation. The UPLC-MS/MS response of prochloraz was evaluated for linearity using a calibration curve of standard solutions in mobile phase A containing 5% matrix and 10 ug/L IS. Concentrations were 0, 0.1, 1.0, 2.5, 5, 7.5, 10 and 15 μg L^−1^. The method limit of detection (LOD) was defined as S/N=3 and estimated from obtained S/N ratio in lowest spiked sample (i.e., 0.1 μg L^−1^). LOQ was extrapolated from LOD multiplying with 10/3.3. Resulting LOD was 0.01 μg L^−1^ and LOQ was 0.03 μg L^−1^.

### D. magna reproduction and length measurements

When treatments were changed the mother was moved with a glass pipette to a new vial and offspring collected and counted. Every 2^nd^ day the following physiological endpoints were recorded: time to first eggs in pouch, time to first clutch, viability of offspring, offspring per clutch, length of offspring in the 2^nd^ clutch, total offspring and total clutches per mother until 5^th^ clutch or until natural death if 5^th^ clutch was not reached. The second clutch was transferred to a transparent slide, which was scanned (Canon CanoScan LiDE 220, 1200dpi, 48 bit colour) and animal length measured according to the method by Agatz and Preuss (2015). When the mothers had delivered their 5^th^ clutch (or at death if before 5^th^ clutch) their length was also measured, and they were subsequently frozen in liquid nitrogen.

### Cytochrome P450 ECOD activity

7-ethoxycoumarin-O-dealkylation (ECOD) activity, which yields the fluorescent product 7-hydroxy coumarin, was used as a measurement of CYP biotransformation capacity of the animals. It is a broad-spectrum substrate for the measurement of overall CYP activity and follows the method by (*30*). Ten animals from the 3rd clutch of each replica was grown to day 5 in a 250 mL blue cap bottle. On day 5 they were transferred to 3 mL glass amber vials and incubated with 7-ethoxycoumarin dissolved in M7 medium and 0.008% acetonitrile (2 mL, 0.02 mM) for 3.5 h at room temperature (25°C). Every 30 min 100 μL aliquots were transferred to a black microwell plate (Greiner Bio-one flat bottom 96-well plate, VWR) and fluorescence was measured (excitation: 380 nm, emission: 480 nm) on a microplate reader (SpectraMax M5 Microplate Reader, Molecular Devices, U.S.) at room temperature. Background contribution to fluorescence was recorded on blank samples containing only incubation medium.

#### Data analysis

After background subtraction, cytochrome P450 activity (change in fluorescence per hour, flu h^−1^) was determined as the slope of a linear regression line fitted in the linear range of fluorescent product formation. Fluorescence was transformed into units of pmol product via calibration standards of 7-hydroxycoumarin in the range 0–140 pmol and was normalized to the length of the animals. Normalization to length rather than protein content was preferred, as this is less time demanding, non-destructive and length and protein constant are proportional until the *Daphnia* starts to produce eggs (>6 days old) (Figure S2).

### Metabolomics

An untargeted metabolomics analysis workflow was applied using two platforms: a nano-liquid chromatography (LC)-Orbitrap HRMS/MS system and an ion exchange chromatography (IC)-orbitrap system. 5-15 animals from the 5^th^ clutch of each replicate mother, were reared to day 5 in a 250 mL blue cap bottle. Subsequently the animals were collected (sieve) and transferred to a cryovial, and immediately submerged in liquid nitrogen and stored at −80°C until extraction.

#### Metabolite extraction

Metabolites were extracted in two fractions for subsequent analysis on the IC- and LC-HRMS platforms. Procedural blanks were included from the beginning. Initially sample homogenization in 1 mL ice-cold methanol/water (1:1) was performed with bead beating under cooling (1 min, 4 m/s, <5 °C) using 1.4 mm ceramic beads and a Bead Ruptor Elite connected to an Omni BR-Cryo cooling unit (Omni International, USA). After centrifugation (15,000 g, 10 min, 4 °C), 500 μL of the supernatant was collected for SPE-cleanup and subsequent LC-HRMS analysis and the remaining supernatant was purified in a liquid-liquid extraction for analysis by IC-HRMS.

#### Liquid-liquid extraction

This procedure was based on the method in Wu et al., (2008) with modifications as in Taylor et al., (2009), but with replacement of chloroform by dichloromethane (DCM). 400 μL DCM was added to the remaining sample and bead beating was repeated (1 min, 4 m/s, <5 °C). After a 10-min incubation on ice to allow for partitioning the samples were centrifuged (15,000 g, 5 min, 4°C). 200 μL of the upper, polar phase was collected for a composite quality control (QC) sample, while another 200 μL aliquot was added directly to a 96-well plate. After vortex mixing the QC was aliquoted into the well plate in equal volume to the samples (200 μL) and the plate was evaporated to dryness in a vacuum concentrator (SpeedVac SPD 1030, Thermo Scientific, Germany) at 35 °C, vacuum of 30 Torr/min down to 5.1 Torr, and reconstituted in 60 μL 50% methanol. The samples were analyzed by IC-HRMS

#### SPE-clean up

250 μL of each sample was combined to make a composite quality control (QC) sample and after a vortex to mix, this pooled sample was aliquoted to eppendorf tubes (V=250 μL). All samples and QCs were put in a vacuum concentrator (SpeedVac SPD 1030, Thermo Scientific, Germany) at 35°C, vacuum of 30 Torr/min down to 5.1 Torr for 1 hr to evaporate methanol. An SPE plate (SOLAμ^TM^ HRP 2mg/1 mL 96 well plate, ThermoFisher Scientific, Germany) was put on a positive pressure manifold and conditioned with 200 μL methanol and 200 μL water with 0.1% formic acid (FA). To load the samples another 200 μL 0.1% FA was added to the reservoirs followed by the samples and QCs and the positive pressure applied. Subsequently the columns were washed with 200 μL 0.1% FA and, after 10 min drying under pressure, eluted into a glass-covered 96-well plate with 2×100 μL methanol. The samples were evaporated to dryness in a vacuum concentrator (SpeedVac SPD 1030, settings as above), and reconstituted in 60 μL 5% methanol and analyzed by nanoLC-HRMS.

#### LC-HRMS/MS

An untargeted analysis workflow was applied using a nanoLC-Orbitrap system. For the LC separation, a Dionex^TM^ UltiMate 3000 RSLCnano system (Thermo Scientific^TM^, Bremen, Germany) was used. The autosampler and columns were thermostated at 8 °C and 40°C, respectively. An amount of 20 μL sample were loaded on a preconcentration trap (C18, 300 μm × 5 mm, 5 μm, 100 Å cartridge) and eluted onto an analytical column (75 μm × 250 mm, 2 μm C18) with a chromatographic triple-phasic 30 minutes gradient delivered at a 300 nL per minute flow rate. The programmed gradient was 10% mobile phase B from 0 to 2 minutes, followed by a gradient reaching 95% B at 17 minutes (curve 5) with an additional 6 minutes plateau (23 minutes) and returning to 10% B at 23.5 minutes. The column was re-equilibrated with 10% B until 30 minutes. The loading solvents A and B were composed of water with 1% and 98% acetonitrile, respectively, and with 0.05% trifluoroacetic acid (TFA). The mobile phases A and B were composed of water with 2% and 98% acetonitrile, respectively, and 0.1% formic acid (FA). The mass spectrometric analysis was performed with a Q Exactive HF Orbitrap (Thermo Scientific, Bremen, Germany) equipped with an EASYspray ion source operated at a spray voltage of 1.80 kV, a capillary temperature at 250 °C, S– lens RF level at 50 V and probe heater temperature at 350 °C. The instrument was operated in data-dependent mode by automatically switching between MS and MS/MS fragmentation. Based on a survey MS scan in the Orbitrap, operated at a mass resolution of 240,000 at m/z 200 with a target of 3e6 ions and a maximum injection time at 100 ms, the 20 most intense ions were selected for MS/MS fragmentation in subsequent scans. The selected ions were isolated at a m/z 0.4 window and higher-energy collision dissociation was done at two normalized collision energies (15 and 50) and fragments recorded in centroid mode at a resolution of 15,000 with a 50 ms max filling time and target of 2e4 ions. An external mass calibration were carried out weekly, using a certified mixture of caffeine and Ultramark 1621 (ThermoFisher Scientific) (*55*).

#### IC-HRMS/MS

An untargeted analysis workflow was applied using a reagent-free ion exchange chromatography system with eluent generation (RFIC-EG) hyphenated with a high-field Orbitrap platform (Thermo Scientific, Bremen, Germany). The autosampler and columns were thermostated at 8 °C and 40°C, respectively. Ten-microliter sample were loaded at a 0.45 mL/minute flowrate onto an anion separator (2 × 250 mm, 4 μm, AS19, Thermo Scientific, Bremen, Germany) with a 50 mm guard with the same separator material. The analytes were passed through a conductivity detector and mixed with LC-MS grade isopropanol (Thermo Fisher Chemicals, Germany) via a tee-piece before infusion into the mass spectrometer. The mass spectrometric analysis was performed with a Q Exactive HF Orbitrap (Thermo Scientific, Bremen, Germany) equipped with an HESI-II ion source (Thermo Scientific, Bremen, Germany) operated at a spray voltage of 3.50 kV, a capillary temperature at 250 °C, S–lens RF level at 50 V and probe heater temperature at 350 °C. The Orbitrap was operated at a mass resolution of 240,000 at m/z 200 with a target of 3e6 ions and a maximum injection time at 100 ms, and the 5 most intense ions were selected for MS/MS fragmentation in subsequent scans. The selected ions were isolated at a m/z 0.4 window and higher-energy collision dissociation was done at 30 NCE and fragments recorded in centroid mode at a resolution of 30,000 with a 100 ms max filling time and target of 1e5 ions.

#### Data analysis

Compound Discoverer software version 3.1.305 (Thermo Scientific) was used for data processing (*56*). The workflow combined the search for unknown metabolites with a search for expected compounds resulting from prochloraz i.e. transformation products and mother compound. The software performed retention time alignment and analyte detection based on the presence of the exact mass of precursor ions, with a mass tolerance of 3.0 ppm. Mass tolerance for spectral alignment was set to 3.0 ppm and a maximum retention time shift of 2.0 min was allowed. The detection of compounds was based on a mass tolerance of 2.0 ppm, intensity threshold of 30%, S/N ratio of 3.0 and a minimum peak intensity of 1,000,000. Compounds were grouped with a mass tolerance of 2.0 ppm and retention time tolerance of 1.0 min. Values below the intensity threshold was marked as gaps but filled with an attempted integration with a mass tolerance of 2.0 ppm and S/N threshold of 1.5. Sample compounds were marked as background if the ratio to a blank sample was below 10. Similarly, the software was set to detect the expected compound prochloraz, dealkylation and dearylation products as well as bio-transformation products with resolution-aware isotope pattern matching. Prediction of molecular composition was done with a mass tolerance of 2.0 ppm, S/N of 3.0, pattern matching intensity threshold of 30% and fragment matching applied. The following maximum element counts: C90, H190, I4, Cl4, N10, O15, P3, S5 were allowed. The compounds marked as background according to the selected criteria in Compound Discoverer were removed, and remaining spectrums visually assessed. The areas of the compounds that fulfilled the quality criteria were exported and normalized to number of animals in the vial. The final number of replicate samples in the analysis were for F1: 6 controls, 6 continuously exposed and 6 F0-exposed and for F2: 9 controls, 8 continuously exposed and 8 F0-exposed and for F2: 9 controls, 8 continuously exposed.

#### Statistical analyses

Quality and validity of the chemical analysis (Figure 2A) was confirmed by principal component analysis (PCA) showing that the composite quality control (QC) samples were centrally located in the plot (Figure 2A). The statistical analyses were performed using R studio v1.2.1335 (R Core Team 2017) and the package DESeq2 v1.22.2 (*57*) to identify statistically significant differences between treatments in relative metabolite concentrations. A variance stabilizing transformation integrated in the DESeq2 package was applied. DESeq2 outputs were also used as input to the principle component analysis and volcano plots and visualized using ggplot2 (Wickham, 2016). DESeq2 uses a negative binomial model to test for significant differences in count data using estimates of variance-mean dependence (*57*). This has been found to have high precision and stable sensitivity in label-free quantative proteomics (*58*) and was therefore found suitable, in spite of the fact that it can be discussed whether the metabolomics data is actually count data.

#### Metabolite identification and functional analyses

Metabolite identification was based on the 4 levels of confidence described in Viant et al. (2019). Putatively annotated compounds (Level 2) was based on comparisons of the experimental MS^2^-spectrum with the online spectral library, mzCloud, and in-house spectral libraries. Putatively characterized compound classes (Level 3) was based on assigned predicted composition and the plausible, tentative candidates to that composition. For instance, whether the compound was likely to naturally occur in *Daphnia* or could be related to prochloraz exposure. Compounds that did not meet these criteria were assigned to level 4 and not incorporated in the functional analysis. The functional analysis was based on the Kyoto Encyclopedia of Genes and Genomes (KEGG) database (Kanehisa et al., 2012). Metabolites of prochloraz (C_15_H_16_N_3_O_2_Cl_3_) were tentatively annotated based on the predicted composition in Compound Discoverer. Increased confidence was achieved by presence of Cl_3_ patterns in the spectrum, compound class scoring, which searched for known fragments of prochloraz, and existing data on invertebrate metabolism of the fungicide (*59*).

### RNA sequencing

10 animals, from the 4th clutch of each F2-mother, were reared to day 5 in a 250 mL blue cap bottle. Subsequently the animals were transferred to a cryovial, liquid was removed and 500 μL RNA-later added. The vials were incubated for 2 hours at 4°C before being transferred to −80°C and stored until RNA extraction.

#### RNA extraction

Total RNA was extracted from 10 animals from the 4^th^ clutch of F2 animals from each replicated treatment line. Total RNA was isolated using TRIzol reagent (ThermoFisher Scientific, USA). The protocol followed manufacturers instruction with modifications as in Campos et al., 2018. In brief, RNAlater was removed, 0.5 mL TRIzol added and homogenization was performed by bead beating using a TissueLyser II (Qiagen, Germany) with 0.5 mm glass beads (Scietific Industries Inc., USA) for 2×45 sec at 30 Hz using cooled beads and 30 sec cooling on dry ice in-between. Insoluble material was removed by centrifugation at 12 000 g (10 min, 4 °C). After transferring the supernatant to a new tube, phenol-chloroform extraction was performed by adding 0.2 mL chloroform per 1 mL of trizol. After shaking by hand, the tubes were left to incubate at room temperature for 2 min before centrifugation at 12 000 g (15 min, 4 °C). Subsequently the aqueous phase was tranferred to a new tube and RNA precipitated with isopropanol (0.5 mL per 1 mL of trizol, VWR, Denmark) by incubation at −20 °C for two hours followed by cenrifugation (12 000 g, 10 min, 4 °C). The pellet was washed with 75% ethanol (1 mL per mL of trizol, centrifugation at 7500g, 5min, 4°C). After re-dissolving in 30 μL DEPC-treated water, DNase treatment was performed by incubation at 37 °C for 30 min using 3 μL DNAse buffer, 1U DNAse (Ambion^TM^ DNAse I, ThermoFisher Scientific, USA) and 10U RNAse inhibitor (Protector RNase inhibitor, Sigma-Aldrich, Germany). Extraction was performed twice with phenol:chloroform:isoamyl alcohol at a ratio of 25:24:1 and once with chloroform. RNA was precipitated overnight at −20°C by addition of 2 volumes of 100% ethanol and 0.1 volume of 3M sodium acetate. After centrifugation (15.000 g, 15 min, 4°C) the RNA was washed with 70% ethanol and resuspended in 30 μL DEPC water. Samples were stored at −80 °C until sequencing.

#### RNA sequencing, assembly and analysis

All library constructions and sequencing were performed at Beijing Genomics Institute (BGI, HongKong). RNA quality was assessed using an Agilent 2100 bioanalyzer and 19 samples (E: n=5, F: n=7, C: n=6) were of sufficient quality to be sequenced using the BGISEQ-500RS sequencing platform. Quality filtering and adaptor removal was performed using Trimmomatic v0.38 (*60*) and quality of reads was evaluated using FastQC (v0.11.5, Babraham bioinformatics). Trimmed reads were mapped to the *D. magna* genome (NCBI database; Assembly: ASM399081v1, BioProject: PRJNA490418), using the alignment software STAR with default settings (*61*). The 123 Mbp *D. magna* genome consists of 4,192 scaffolds and 16,817 contigs, with 15,721 annotated genes (*62*). The raw counts of the number of mapped transcripts were obtained with FeatureCounts (*63*), and the coverage uniformity and transcript integrity was assessed using the RSeQC package (*64*). This revealed a substantial 3’ bias (Figure S3), and consequently 4 replicate samples that showed considerable degradation were removed. Normalization of the remaining samples was performed using the transcript integrity number (TIN) as described in (*65*). Finally the R-package DESeq2 v1.22.2 (*57*), R Core Team 2017), was used to identify differentially expressed genes. During data analysis it became evident that the physical length of *D. magna* was a key predicting variable of variation in gene expression between replicate samples. It was not possible to measure the length of the animals used for RNA extraction as the increased stress of the measuring procedure would almost certainly have affected sample transcription levels. Length measurements of daphnids from the ECOD analysis of the previous clutch was therefore used as a proxy for the length of animals in the transcriptomic analysis. Unfortunately, one length-measurement was missing and the final number of replicate samples in the analysis after filtering and normalization were therefore: 5 controls, 4 continuously exposed and 5 F0-exposed, which each consisted of RNA-extracts from 8 five-day old *D. magna* juveniles.

### Statistics

Data for time to first clutch and ECOD-activity were modelled by analysis of covariance (ANCOVA) with generation and treatment as explanatory variables. In the analysis of length, amount and mortality of offspring a linear mixed model of the R-package “lme4” (*66*) was applied. In all cases the initial model was an additive model consisting of the two explanatory variables. Subsequently, interactions between the explanatory variables were tested and if interactions were found the co-variates were combined into one explanatory variable, which was used in the final model. An interaction between treatment and generation was found for all models. The linear mixed models for neonate length included the replicate as a random variable. The linear mixed model for neonate amount and mortality was adjusted for the day of measurement and the clutch. Model checking was performed by visual assessment of QQ-plots and residual plots and if okay, a *post-hoc* pair-wise comparison by t-tests (Tukey contrasts) was performed (Hothorn *et al.*, 2008). For each analysis a pre-specified significance level of 5 % was used. All analyses were carried out using R v3.6.3 and R studio v1.1.383 (R Core Team 2017).

## Acknowledgements

We thank research fellows Xiaogang Jiang, Amandine Levastre, Michele Gottardi and Anja Weibel for help with the laboratory work, Andrea Rösch for sharing her knowledge on prochloraz metabolism in invertebrates, Bruno Campos for advice on RNA extractions from *Daphnia*, Arnar Pálsson and Dagný Ásta Rúnarsdóttir for assistance and academic advices in data treatment of RNAseq data.

## Author Contributions

R.P. and N.C. conceived and designed all the experimental work. R.P fabricated samples. H.H.D.F.L contributed to the transcriptome methodology and assisted with the data analysis. M.H. contributed to the metabolome methodology and assisted the measurement and characterization. The manuscript was written by R.P. All authors contributed to the discussion of the data, proofread of the manuscript. The overall project was supervised by N.C. and H.H.D.F.L.

## Competing interests

The authors declare no competing interests.

## Data availability

All data needed to assess the conclusions of the paper are given in the main text, supplementary material and data file S1. Metabolomics data is furthermore available in Metabolights under study accession number MTBLS1836 and RNAseq data is available through ArrayExpress under study accession number E-MTAB-9309

## Funding

The authors acknowledge financial support from University of Copenhagen, Department of Plant and Environmental Sciences, PhD scholarship for R. Poulsen and M. Hansen acknowledge the financial starting grant from Aarhus University Research Foundation (AUFF-T-2017-FLS-7-4). The Villum Foundation (Young Investigator Grant, grant number 10122 to HHDFL), and a The Independent Research Fund Denmark Sapere Aude Research grant no 8049-00086B to HHDFL

## Notes

### Competing Interest Statement

The authors have declared no competing interest.

