## Supplementary information for "Grandmother’s chemical legacy of pesticide exposure: bi-generational effects and acclimation in a model invertebrate"

**This PDF file includes:**

Supplementary text
Figures S1 to S5
Tables S1 to S9

**Other supplementary materials for this manuscript include the following:**

Dataset S1

### Pilot study results

A 21-day pilot study, based on the OECD guideline for *Daphnia magna* reproduction test (OECD/OCDE, 2008) was performed with the concentration range 0, 10, 50, 100, 250 and 500 μg/L prochloraz. Mothers were kept individually in 100 mL beakers containing 50 mL test solution, and there were 15 replicate glasses for each concentration. Glasses were checked every 24 h for health status of mothers and the medium and treatment was renewed every 3 days. On change days, offspring were collected, counted, scanned and their length measured. Furthermore, the cytochrome P450 ECOD activity of mothers was measured at termination (day 21). Figure S1 sums up the dose-response relationship for the endpoints of cumulative offspring per mother, length of offspring and cytochrome P450 ECOD activity in mothers.

The data was fitted with a 4-parameter log-logistic dose-response model (LL4 in the R package drc):

$$y = c + \frac{d-c}{1+\exp\,[b((\log(x)-\log(e))]} \qquad \text{Eq. S1}$$

Where c is the lower limit, d is the upper limit, e is the concentration giving 50% response ($EC_{50}$) and b is the slope of the curve around the $EC_{50}$- value. When appropriate the model was reduced to a 3-parameter model where the lower limit (c) is set to 0 (LL3). Parameters for the curve fits are given in Table S1

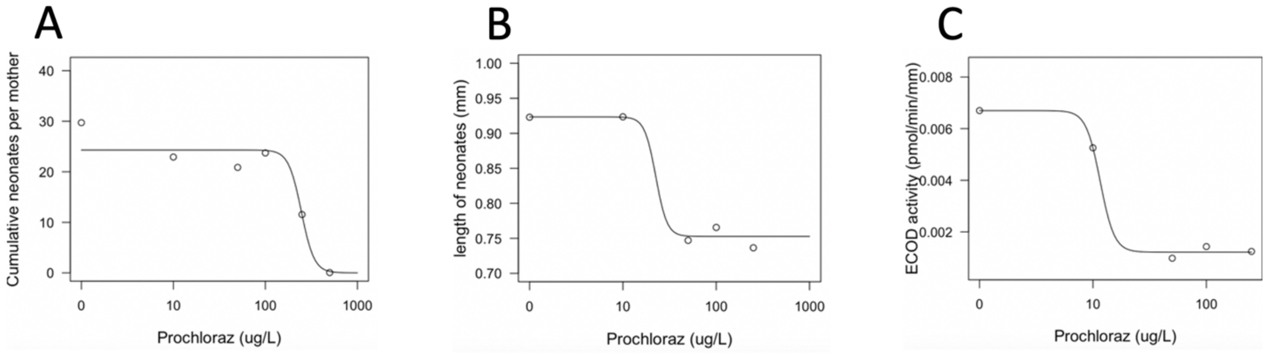

**Figure S1**. A 21-day pilot study, based on the OECD guideline for *Daphnia magna* reproduction test (OECD/OCDE, 2008) was performed with the concentration range 0, 10, 50, 100, 250 and 500 μg/L prochloraz to investigate dose-response relationships for **(A**) Cumulative number of offspring (neonates) per mother at day 21 **(B)** length of offspring **(C)** Cytochrome P450 ECOD activity in mothers at day 21. Dots represents the average of replicates (*n* = 15) and data was fitted by log-logistic dose-response models (R package drc) and the parameters for each model can be found in Table S1

**Table S1.** Data from the pilot study was fitted by log-logistic dose-response models (R package drc). Values for estimated model parameters ± standard error of the estimate as well as the type of model and the residual standard error for the fit is provided.

| | model | b | c | d | e | Residual SE |
|---|---|---|---|---|---|---|
| Cumulative no. of offspring | LL3 | 6 ± 11 | NA | 24±1 | 246±18 | 7.2 (80 DF) |
| Length of offspring | LL4 | 7±12 | 0.752±0.008 | 0.92±0.01 | 23± 32 | 0.22 (1301 DF) |
| ECOD activity in mothers | LL4 | 7±44 | 1.2e-3 ±3e-04 | 6.7e-03±5e-04 | 11.6 ± 11 | 0.00086 (10 DF) |

**Feeding scheme**

Animals were fed with green algae (*Raphidocelis subcapitata*), re-suspended in newly-prepared M7 medium (A(684 nm) = 0.5 corresponding to approximately $4.2*10^5$ cells/mL). Feeding was done as part of the treatment change: when the new treatment was prepared, algae suspension was added as part of the 500 mL treatment. The feeding scheme was as follows:

- Day 0 and 2: 12.5 mL
- Day 4 and 6: 20 mL
- Day 8-22: 30 mL
- > Day 21: 35 mL

Feeding of the growing juveniles were also only done initially; when the new treatment was prepared, 6.25 mL algae suspension was added as part of the 250 mL treatment. The feeding scheme resulted in *ad libitum* food for all animals as algae were always present at the bottom of exposure vessels.

### Correlation between length and protein content in *D. magna*

Figure S2 shows the correlation between protein content and length in *D. magna*. The linear regression coefficient is 0.9337. The length of animals was measured as specified in Materials and Methods. The protein content was measured according to the Bradford protein assay based on the original Bradford method (Bradford, 1976). Animals were homogenized in phosphate buffer (0.3 M, pH 7.5) using an ultrasonic sonicator with a cycle of 3 x 3s and amplitude of 20 %. 5µL of the homogenates was incubated with 250 µL Bradford reagent (Sigma Aldrich, Germany) at room temperature for 10 min in a transparent micro well plate. Absorbance was measured at 595 nm on a microplate reader (SpectraMax M5 Microplate Reader, Molecular Devices, U.S.) at room temperature. Protein content was quantified using a bovine serum albumin (BSA) standard curve (0, 0.1, 0.25, 0.5, 0.75, 1, 1.2, 1.4 mg prot/mL in phosphate buffer 0.3 M, pH = 7.5).

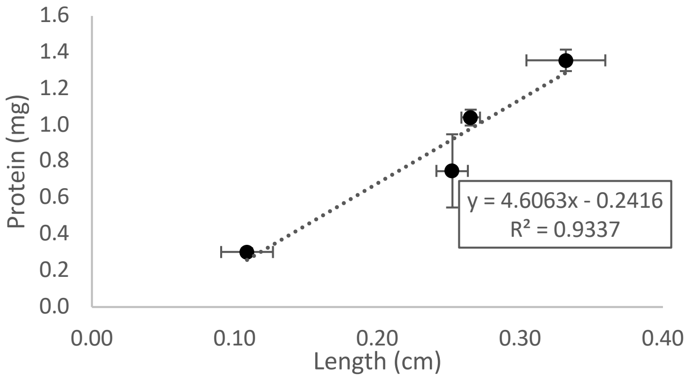

**Figure S2.** Protein content (mg) of *Daphnia magna* as a function of length fitted with a linear regression model

**Bias in coverage in the RNAseq data**

Coverage uniformity and transcript integrity was assessed using the RSeQC package (Wang et al., 2012). The function 'geneBody_coverage.py' scales all the transcripts to 100 nucleotides (nt) and calculates number of reads that cover each nucleotide position. A plot can be generated to visualize the coverage uniformity over the gene body (Figure S3). When the transcription of a gene terminates, polyadenylation (addition of a poly-A tail) happens at the 3' end of the transcript and this part is therefore better protected from degradation. Hence, the 5' is more likely to be degraded crating a bias in the coverage.

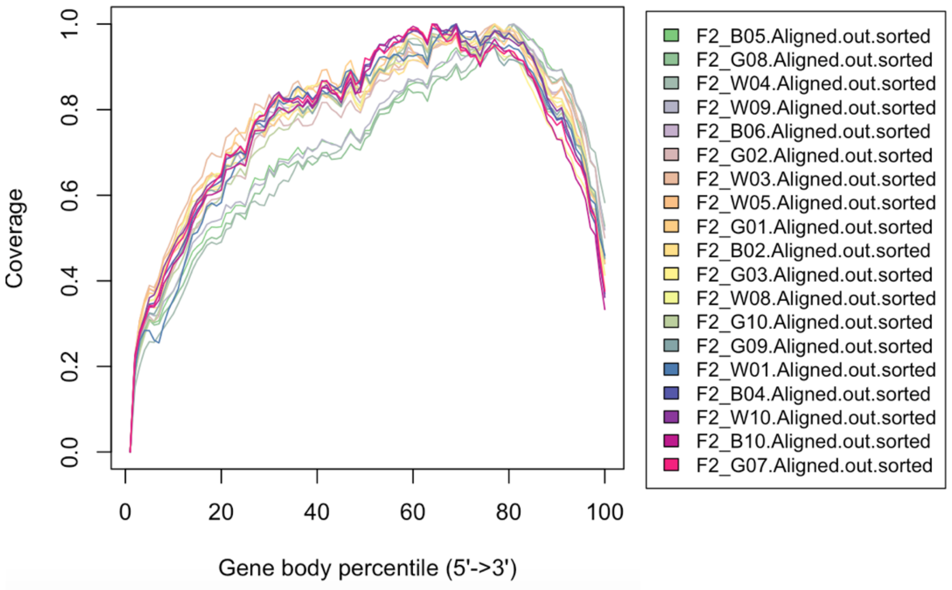

**Figure S3.** Relative number of reads (coverage) over the gene body plotted in the 5'→3' direction to visualize coverage uniformity in the RNA sequencing data. Using the RSeQC package, all transcripts were scaled into 100 nucleotides (nt) and number of reads that cover each nucleotide position calculated.

**Exposure verification**

Water samples were collected from three random vials within each treatment group before change and from each of the new test solutions before distribution into test vials. Prochloraz concentration was quantified using UHPLC-MS$^2$ with deuterated prochloraz (d4-prz) as internal standard. Figure S4 shows the measured concentrations over the time course of the experiment. In continuously exposed animals (Figure S4.A) the average start concentration was 87±10 µg/L and this decreased with 7% during the 48 hours that passed between treatment changes. For F0-exposed animals (Figure S4.B), the average start concentration in F0 was 71 ± 25 µg/L and this decreased 62 ± 22 µg/L over 48 hours. Problems with pump pressure on the instrument during these measurements is the most likely reason for the larger standard deviations. For generation F1 and F2, where no exposure should happen, 10 randomly selected samples were measured from each generation. No concentrations of prochloraz above LOD (0.01 µg/L) was detected. Also, in control animals (FigureS4.C) 10 samples per generation were randomly selected for measurement and no traces of prochloraz was found.

Table S2 and S3 shows the average concentrations of prochloraz in growing bottles over the 5 days. Only the rearing bottle of treatment ”Exposed” should contain prochloraz. The average start concentration was 71±5 µg/L and this decreased with 3% during the 5 days that passed between treatment changes. Unfortunately, a mistake was made in the preparation of the exposure-media for the F1 animals of treatment E used for ECOD activity measurements (except for replicate E07): Hence, these were only subjected to 37.5 µg/L during the rearing phase from offspring to 5 days age. Prochloraz was not detected above LOD in 10 water samples random selected from F0-exposed and Control treatments. Special care was taken to measure all samples from the F0-exposed treatment group, where animals were moved from a prochloraz containing solution to pure medium, to make sure that no significant carry-over had happened. This was not the case.

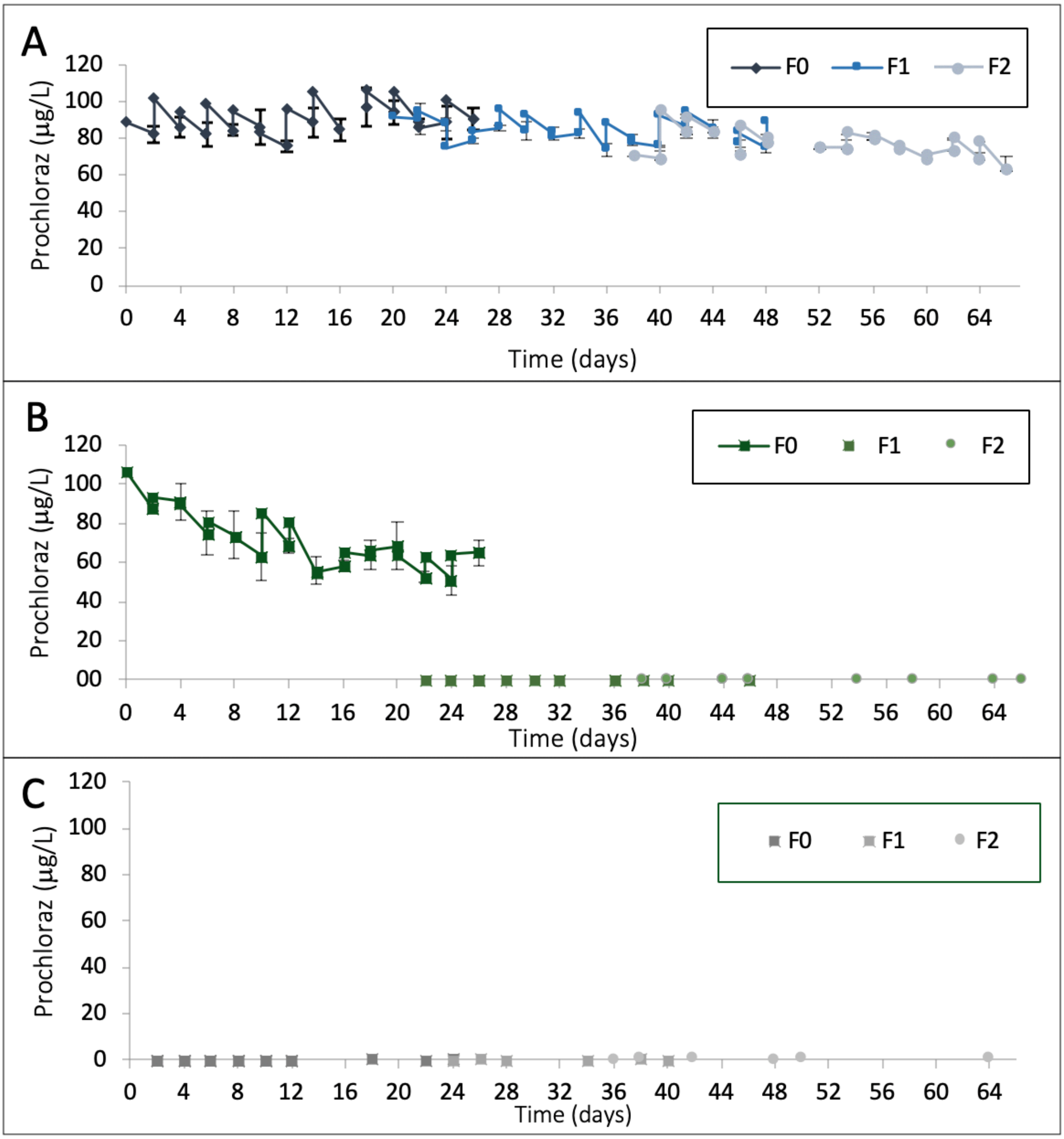

**Figure S4.** Exposure scenario for **(A)** continuously exposed animals, where the average initial concentration was 87±10 µg/L, which decreased with 7% on average to 81 ± 8 µg/L over 48 hours**. (B)** F0-exposed animals, where average initial concentration in F0 was 71±25 µg/L, which decreased to 62 ± 22 µg/L over 48 hours and all samples from F1 and F2 were < LOD of 0.01 µg $L^{-1}$ **(C)** non-exposed (control) animals, where all measured samples were < LOD of 0.01 µg $L^{-1}$. Error bars show standard deviation (n=3).

**Table S2.** Results for exposure verification in generation F1 and the three treatments: Exposed, F0-exposed and control. All replicates are listed along with measured concentration on day 1 and day 5, if performed. LOD for the method was 0.01 μg $L^{-1}$ and LOQ was 0.03 μg $L^{-1}$

| Exposed | Day 1 | Day 5 | F0-exposed | Day 1 | Day 5 | Control | Day 1 | Day 5 |
|---|---|---|---|---|---|---|---|---|
| F1-ECOD-E-a | 37.5 | | F1-ECOD-F-a | 0.0 | | F1-ECOD-C-a | | |
| F1-ECOD-E-b | | | F1-ECOD-F-b | 0.0 | | F1-ECOD-C-b | 0.0 | 0.0 |
| F1-ECOD-E-c | 37.5 | | F1-ECOD-F-c | 0.0 | | F1-ECOD-C-c | | |
| F1-ECOD-E-d | 37.5 | 34.7 | F1-ECOD-F-f | 0.0 | | F1-ECOD-C-d | | |
| F1-ECOD-E-e | 37.5 | 34.3 | F1-ECOD-F-g | 0.0 | | F1-ECOD-C-e | | |
| F1-ECOD-E-f | 37.5 | 32.3 | F1-ECOD-F-h | 0.0 | | F1-ECOD-C-f | | |
| F1-ECOD-E-g | 72.6 | | F1-ECOD-F-i | 0.0 | | F1-ECOD-C-g | | |
| F1-ECOD-E-h | 37.5 | 75.3 | F1-ECOD-F-j | 0.0 | | F1-ECOD-C-h | 0.0 | 0.0 |
| F1-ECOD-E-i | 37.5 | 32.8 | | | | F1-ECOD-C-i | | |
| F1-ECOD-E-j | | | | | | F1-ECOD-C-j | | |
| F1-RNA-E-a | 64.2 | 58.3 | F1-RNA-F-a | 0.0 | | F1-RNA-C-a | | |
| F1-RNA-E-b | 70.0 | 72.1 | F1-RNA-F-b | 0.0 | | F1-RNA-C-b | | |
| F1-RNA-E-c | 37.5 | 36.1 | F1-RNA-F-c | 0.0 | | F1-RNA-C-c | | |
| F1-RNA-E-d | 37.5 | 33.7 | F1-RNA-F-f | 0.0 | | F1-RNA-C-d | | |
| F1-RNA-E-e | | | F1-RNA-F-g | 0.0 | | F1-RNA-C-e | | |
| F1-RNA-E-f | 73.5 | 73.4 | F1-RNA-F-h | 0.0 | | F1-RNA-C-f | | |
| F1-RNA-E-g | | | F1-RNA-F-i | 0.0 | | F1-RNA-C-g | | |
| F1-RNA-E-h | 69.6 | 65.3 | F1-RNA-F-j | 0.0 | | F1-RNA-C-h | | |
| F1-RNA-E-i | 65.7 | 66.5 | | | | F1-RNA-C-i | 0.0 | 0.1 |
| F1-RNA-E-j | | | | | | F1-RNA-C-j | | |

**Table S3.** Results for exposure verification in generation F2 and the three treatments: Exposed, F0-exposed and control. All replicates are listed along with measured concentration on day 1 and day 5, if performed. LOD for the method was 0.01 $\mu g\ L^{-1}$ and LOQ was 0.03 $\mu g\ L^{-1}$

| **Exposed** | **Day 1** | **Day 5** | **F0-exposed** | **Day 1** | **Day 5** | **Control** | **Day 1** | **Day 5** |
|---|---|---|---|---|---|---|---|---|
| F2-ECOD-E-a | | | F2-ECOD-F-a | | | F2-ECOD-C-a | | |
| F2-ECOD-E-b | | | F2-ECOD-F-b | | | F2-ECOD-C-b | | |
| F2-ECOD-E-c | | | F2-ECOD-F-c | | | F2-ECOD-C-c | 0.0 | 0.0 |
| F2-ECOD-E-d | | | F2-ECOD-F-f | | | F2-ECOD-C-d | | |
| F2-ECOD-E-e | | | F2-ECOD-F-g | | | F2-ECOD-C-e | 0.0 | 0.0 |
| F2-ECOD-E-f | | | F2-ECOD-F-h | | | F2-ECOD-C-f | | |
| F2-ECOD-E-g | | | F2-ECOD-F-i | | | F2-ECOD-C-g | | |
| F2-ECOD-E-h | | | F2-ECOD-F-j | 0.1 | 0.0 | F2-ECOD-C-h | | |
| F2-ECOD-E-i | | | | | | F2-ECOD-C-i | | |
| F2-ECOD-E-j | | | | | | F2-ECOD-C-j | | |
| F2-RNA-E-a | | | F2-RNA-F-a | 0.0 | 0.0 | F2-RNA-C-a | | |
| F2-RNA-E-b | 82.0 | | F2-RNA-F-b | 0.0 | 0.0 | F2-RNA-C-b | | |
| F2-RNA-E-c | 72.2 | | F2-RNA-F-c | | | F2-RNA-C-c | 0.0 | |
| F2-RNA-E-d | 70.3 | 66.0 | F2-RNA-F-f | | | F2-RNA-C-d | | |
| F2-RNA-E-e | | | F2-RNA-F-g | | | F2-RNA-C-e | | |
| F2-RNA-E-f | 66.8 | | F2-RNA-F-h | | | F2-RNA-C-f | 0.0 | |
| F2-RNA-E-g | | 79.3 | F2-RNA-F-i | | | F2-RNA-C-g | | |
| F2-RNA-E-h | | | F2-RNA-F-j | | | F2-RNA-C-h | | |
| F2-RNA-E-i | | | | | | F2-RNA-C-i | | |
| F2-RNA-E-j | | | | | | F2-RNA-C-j | | |

**Survival of offspring**

Reproduction was assessed as cumulative number of live neonates as this better reflects the actual effect on the population. Figure S4, however, shows the mortality of offspring as a perspective on this.

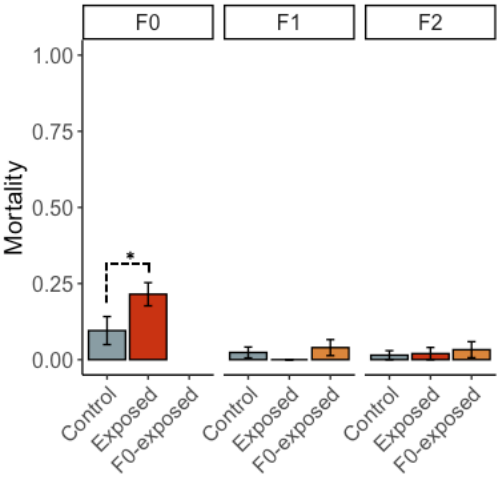

**Figure S5.** Mortality of offspring for the three treatment groups: control (blue), continuously exposed (red) and F0-exposed (orange) in generation F0-F2. Error bars show standard error of the mean (SEM), * indicate significant difference (Tukey´s post hoc *T-test* $p<0.05$).

**Physiological data of control animals**

Offspring of control animals were significantly smaller in F1 compared to F0 and F2, which seems to be a random effect of fluctuating conditions in the test system. The cumulative reproduction in controls furthermore decreased significantly from F0 to F1 and from F1 to F2, while there were no significant differences between generations for the continuously exposed and F0-exposed animals. This decrease in control reproduction could be due to non-optimal housing of the animals in terms of enclosure height and diameter, isolation from other individuals, microbiological changes, etc. (Ebert, 2005; Mucklow and Ebert, 2003; Sison-Mangus et al., 2015). Decreased length of *D. magna* has also been found in cases where food levels were too high compared to the energy consumption of the animals (Porter et al., 1983).

**Table S4.** Functionally annotated metabolites arranged according to KEGG compound classes. For each metabolite the name, KEGG ID and pathway IDs are shown. Furthermore, the confidence level of the ID, the molecular weight (Mw), the formula for the predicted composition as well as the pattern coverage (%) and delta mass of that composition is shown. Finally, the log2 fold change for the comparisons: Continuously exposed versus controls in F1 (F1E vs F1C) and F2 (F2E vs F2C), and F0-exposed versus controls in F1 (F1F vs F1C) and F2 (F2F vs F2C).

| Compound Name | Pathway IDs | KEGG ID | ID-level | Mw | Formula | Platform | Pattern coverage [1](%) | DeltaMass [2] [ppm] | log2 FC: F1E vs F1C | log2 FC: F1F vs F2C | log2 FC: F2E vs F2C | log2 FC: F2F vs F2C | notes |
|---|---|---|---|---|---|---|---|---|---|---|---|---|---|
| **KEGG Class: Biosynthesis of amino acids** | | | | | | | | | | | | | |
| 2-Oxobutyric acid | map01230, 00290, 00250, 00310, 00260 | C00109 | 2 | 102.03159 | C4 H6 O3 | IC | 99.89 | -1.05 | 2.2 | | | | |
| 3-Isopropylmalic acid | map01230, 00290, | C02504 | 2 | 176.06842 | C7 H12 O5 | IC | 98.72 | -0.31 | 1.7 | | | | |
| Adipic acid | map01230, 00290, | C06104 | 2 | 146.05787 | C6 H10 O4 | IC | 99.86 | -0.28 | 1.7 | | | | |
| Citric acid | map01230 | C00158 | 2 | 192.02699 | C6 H8 O7 | IC | 100 | -0.04 | 1.1 | | | | |
| Glutaric acid | map00310 | C00489 | 2 | 132.04221 | C5 H8 O4 | IC | 99.89 | -0.36 | 2.5 | | | | |
| Fumaric acid | map00250, 00360 | D02308 | 2 | 116.01093 | C4 H4 O4 | IC | 100 | -0.24 | 2.2 | | | | |
| N-Acetylvaline | map00310 | | 2 | 159.0895 | C7 H13 N O3 | IC | 99.2 | -0.27 | 1.1 | | | | |
| Malonic acid | map00260 | C00383 | 2 | 104.01094 | C3 H4 O4 | IC | 99.85 | -0.22 | 1.7 | | | | |
| Benzoic acid | map00360 | C00180 | 2 | 122.03672 | C7 H6 O2 | IC | 100 | -0.47 | 2.7 | | | | |
| 4-Hydroxybenzaldehyde | map00360 | C00633 | 2 | 122.03672 | C7 H6 O2 | IC | 100 | -0.46 | 2.4 | | | | |
| **KEGG Class: Metabolism of other amino acids** | | | | | | | | | | | | | |
| Taurine | map00430 | C00245 | 2 | 125.01459 | C2 H7 N O3 S | IC | 99.89 | -0.61 | -0.9 | | | | |
| 2-Hydroxyethanesulfonate | map00430 | C05123 | 3 | 125.99861 | C2 H6 O4 S | IC | 100 | -0.52 | 3.5 | | -2.7 | -7.4 | |
| **KEGG Class: Carbohydrate metabolism** | | | | | | | | | | | | | |
| Citric acid | map01200, 00053, 00630, 00020 | C00158 | 2 | 192.02699 | C6 H8 O7 | IC | 100 | -0.04 | 1.1 | | | | |
| L-Threonic acid | map00053 | C01620 | 2 | 136.03716 | C4 H8 O5 | IC | 99.82 | -0.12 | 1.0 | | | | |
| Malonic acid | map01200, 00053, 00630 | C00383 | 2 | 104.01094 | C3 H4 O4 | IC | 99.85 | -0.22 | 1.7 | | | | |
| δ-Gluconic acid δ-lactone | map01200, 00030, 00053 | C00198 | 2 | 178.04772 | C6 H10 O6 | IC | 98.6 | -0.1 | 2.4 | | | | |
| 3-Isopropylmalic acid | map01200, 00620, | C02504 | 2 | 176.06842 | C7 H12 O5 | IC | 98.72 | -0.31 | 1.7 | | | | |
| Fumaric acid | map00620, 00020, 01200 | D02308 | 2 | 116.01093 | C4 H4 O4 | IC | 100 | -0.24 | 2.2 | | | | |
| 2-Oxobutyric acid | map01200 | C00109 | 2 | 102.03159 | C4 H6 O3 | IC | 99.89 | -1.05 | 2.2 | | | | |

[1]Pattern coverage (%) displays the summed intensity of the matching isotope peaks in the measured MS1 spectrum relative to the summed intensity in the theoretical isotope pattern, thereby giving a quantitation of how well the measured isotope pattern matches the theoretical pattern.

[2]Delta mass (ppm) is the difference between the measured Mw and the theoretical Mw for the predicted composition

**Table S5.** Functionally annotated metabolites arranged according to KEGG compound classes. For each metabolite the name, KEGG ID and pathway IDs are shown. Furthermore, the confidence level of the ID, the molecular weight (Mw), the formula for the predicted composition as well as the pattern coverage (%) and delta mass of that composition is shown. Finally, the log2 fold change for the comparisons: Continuously exposed versus controls in F1 (F1E vs F1C) and F2 (F2E vs F2C), and F0-exposed versus controls in F1 (F1F vs F1C) and F2 (F2F vs F2C).

| Compound Name | Pathway IDs | KEGG ID | ID-level | Mw | Formula | Platform | Pattern coverage [1](%) | DeltaMass [2] [ppm] | log2 FC: F1E vs F1C | log2 FC: F1F vs F2C | log2 FC: F2E vs F2C | log2 FC: F2F vs F2C | notes |
|---|---|---|---|---|---|---|---|---|---|---|---|---|---|
| **KEGG Class: Lipid metabolism** | | | | | | | | | | | | | |
| Caprylic acid | map00061 | C06423 | 2 | 144.11493 | C8 H16 O2 | IC | 100 | -0.67 | 1.4 | | | | |
| Malonic acid | map00061 | C00383 | 2 | 104.01094 | C3 H4 O4 | IC | 99.85 | -0.22 | 1.7 | | | | |
| glutaric acid | map00071 | C00489 | 2 | 132.04221 | C5 H8 O4 | IC | 99.89 | -0.36 | 2.5 | | | | |
| ***alpha-linoleic acid metabolism*** | | | | | | | | | | | | | |
| α-Linolenic acid | map00592 | C06427 | 2 | 278.22434 | C18 H30 O2 | LC | 99.29 | -0.88 | 1.5 | | | | |
| 9(S)-HPOT | map00529 | C16321 | 3 | 310.21416 | C18 H30 O4 | LC | 97.36 | -0.82 | 2.1 | | | | CYP substrate |
| 13(S)-HPOT | map00529 | C04785 | 3 | 310.21414 | C18 H30 O4 | LC | 97.36 | -0.87 | 2.1 | | | | CYP substrate |
| 9,10-EOTrE | map00529 | C16324 | 3 | 292.2037 | C18 H28 O3 | LC | 97.57 | -0.85 | -1.6 | | | | CYP product |
| ***Cutin, suberine and wax biosynthesis*** | | | | | | | | | | | | | |
| Hexadecanedioate | map00073 | C19615 | 3 | 286.21431 | C16 H30 O4 | LC | 85.3 | -0.35 | 2.3 | | | | |
| 10,16-Dihydroxyhexadecanoic acid | map00073 | C08285 | 3 | 288.22994 | C16 H32 O4 | LC | 97.71 | -0.43 | 1.5 | | | | CYP product |
| ***steroid hormone biosynthesis*** | | | | | | | | | | | | | |
| Estriol | map00140 | C05141 | 3 | 288.17234 | C18 H24 O3 | LC | 99.58 | -0.70 | 1.3 | | | | CYP product |
| Tetrahydrocorticosterone | map00140 | C05476 | 3 | 350.24525 | C19 H36 O4 | LC | 98.74 | -1.32 | 2.2 | | -1.8 | | |
| Tetrahydrocortisol | map00140 | C05472 | 3 | 366.24025 | C21 H34 O5 | LC | 98.86 | -1.02 | 2.8 | | | 1.8 | oxidoreductase substrate and product |
| **KEGG Class: Metabolism of cofactors and vitamins** | | | | | | | | | | | | | |
| All-trans-4-Oxoretinoic acid | map00830 | C16678 | 3 | 314.18796 | C20 H26 O3 | LC | 99.03 | -0.75 | | -2.2 | | | CYP product |
| **KEGG Class: Metabolism of terpenoids and polyketides** | | | | | | | | | | | | | |
| ***Sesquiterpenoid and triterpenoid biosynthesis*** | | | | | | | | | | | | | |
| Costunolide | map00909 | C09382 | 3 | 232.14628 | C15 H20 O2 | LC | 98.41 | -0.22 | 1.2 | | | | CYP product |
| Germacrene A acid | map00909 | C19678 | 3 | 234.16185 | C15 H22 O2 | LC | 99.43 | -0.54 | -1.3 | | | | CYP substrate |
| ***Diterpenoid biosynthesis*** | | | | | | | | | | | | | |
| 6beta,7beta-Dihydroxykaurenoic acid | map00904 | C11876 | 3 | 334.21422 | C20 H30 O4 | LC | 86.1 | -2.62 | 1.5 | -4.7 | | | . |
| ***Ubiquinone and other terpenoid-quinone biosynthesis*** | | | | | | | | | | | | | |
| Geranylhydroquinone | map00130 | C10793 | 3 | 246.16191 | C16 H22 O2 | LC | 98.26 | -0.28 | 1.9 | | | | |
| ***Insect hormone biosynthesis*** | | | | | | | | | | | | | |
| juvenile hormone III acid | map00981 | C16504 | 3 | 252.17239 | C15 H24 O3 | LC | 99.43 | -0.63 | 1.5 | | -1.5 | -1.1 | CYP product |

**Table S6.** Prochloraz metabolites tentatively identified (level 2). For each metabolite an assigned ID number, the retention time (RT), the molecular weight (Mw), the formula for the predicted composition as well as the pattern coverage (%) and delta mass (ppm) of that composition is shown. Additionally, the log2 fold change for the comparisons: Continuously exposed versus controls in F1 (F1E vs F1C) and F2 (F2E vs F2C), is shown. Finally, the confidence level of annotation, the suggested transformation from the parent compound, the composition change and a suggested structure when possible, is given.

| ID | RT (min) | Molecular weight | Formula | Pattern coverage (%) | DeltaMass [ppm] | log2 FC: F1E vs F1C | log2 FC: F2E vs F2C | ID-level | Transformation | Composition change | Suggested Structure |
|---|---|---|---|---|---|---|---|---|---|---|---|
| Prochloraz (LC.27) | 17.526 | 375.03063 | C15 H16 Cl3 N3 O2 | 98.7 | -0.43 | 11.1 | 8.4 | 2 | Parent compound | | 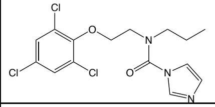 |
| LC.768 | 19.801 | 352.01469 | C13 H15 Cl3 N2 O3 | 95.56 | -0.3 | 10.6 | 8.5 | 2 | Partial loss of imidazole ring, aldehyde formation* | -C2HN +O | 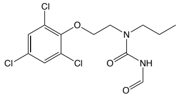 |
| LC.790 | 18.896 | 373.01503 | C15 H14 Cl3 N3 O2 | 95.79 | | 10.3 | 7.1 | 2 | Desaturation | -H2 | 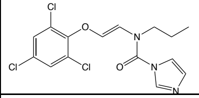 |
| LC.23 | 17.545 | 325.00393 | C12 H14 Cl3 N O3 | 98.83 | -1.13 | 11.3 | 5.3 | 2 | Loss of imidazole ring, Oxidation | -C3H2N2 +O | 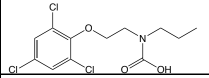 |
| LC.24 | 17.396 | 306.99286 | C12 H12 Cl3 N O2 | 93.79 | -1.54 | 10.9 | 4.9 | 2 | Loss of imidazole ring, desaturation | -C3H2N2 -H2 | 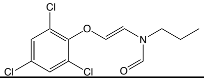 |
| LC.185 | 17.523 | 221.94031 | C8 H5 Cl3 O | 97.07 | -1.3 | 8.3 | 5.8 | 2 | Desaturation, remaining chlorophenyl moiety | -C7H9N3O -H2 | 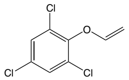 |
| LC.714 | 17.548 | 585.20279 | C26 H42 Cl3 N O7 | 96.37 | 0.17 | 7.7 | 6.6 | 2 | | + C11 H26 O5 -N2 | |
| LC.638 | 17.436 | 573.15526 | C26 H34 Cl3 N3 O5 | 97.44 | -1.99 | 5.9 | 6.0 | 2 | | + C11 H18 O3 | |
| LC.701 | 17.482 | 627.21367 | C28 H44 Cl3 N O8 | 97.42 | 0.75 | 10.6 | 5.2 | 2 | | + C13 H28 O6 -N2 | |
| LC.710 | 17.439 | 629.18152 | C29 H38 Cl3 N3 O6 | 95.27 | -2.3 | 9.1 | 4.3 | 2 | | + C14 H28 O4 | |
| LC.728 | 13.649 | 385.96561 | C12 H13 Cl3 N2 O4 S | 91.01 | -1.44 | 8.9 | 7.7 | | | -C3 H3 N +SO2 | |
| LC.735 | 18.562 | 408.99185 | C12 H18 Cl3 N O4 S | 89.35 | -0.35 | 9.3 | 6.5 | 2 | | -C3H3N2 + H5SO4 | |
| LC.736 | 19.068 | 562.22428 | C28 H45 Cl3 N2 O3 | 91.48 | 2.5 | 11.0 | 9.1 | 2 | Partial loss of imidazole ring, palmitoyl conjugation | + C13 H29 O -N | 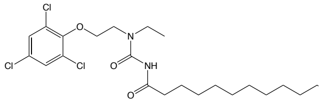 |

[1]Pattern coverage (%) displays the summed intensity of the matching isotope peaks in the measured MS1 spectrum relative to the summed intensity in the theoretical isotope pattern, thereby giving a quantitation of how well the measured isotope pattern matches the theoretical pattern.

[2]Delta mass (ppm) is the difference between the measured Mw and the theoretical Mw for the predicted composition

**Significantly depleted and enriched GO-terms in grand-maternally exposed animals**

Significantly enriched GO-terms (Table S6) are discussed in the main manuscript. Significantly depleted GO-terms were related to the spliceosomal complex, RNA binding and mitochondrial transcription (Table S7). The spliceosome is the molecular machinery that converts precursor-mRNAs (pre-mRNAs) to mRNAs by removing non-coding sequences (intron) and tie together the coding sequences (exons). This enables alternative splicing where different mRNAs can be generated from the same pre-mRNA. The spliceosome is a dynamic complex of several subunits, which can come together in different conformations and composition (Will and Luhrmann, 2011). This gives it an important role in regulatory plasticity (Filichkin et al., 2015; Matera and Wang, 2014), however, atypical splicing have also been connected with pathological outcomes, mainly in humans (Chen and Moore, 2014). No specific conclusions about the consequences of depletion within this gene ontology can be drawn, however, it is clear that the transcriptional landscape is changed in the animals with grandmaternal exposure compared to control animals. The differences in mitochondrial transcription suggests that the grandmaternal exposure to prochloraz had an effect on energy conversion and production, which might be further related to the observed effect on reproductive output (Figure1D), and (Campos et al. (2012)).

**Table S7.** Significantly enriched biochemical pathways identified by their gene ontology (GO) terms in F0-exposed animals compared to controls. The number of significantly up-regulated (DE) and the total number of genes (Total) with this GO-term assigned, as well as the Bonferroni-adjusted p-value for the hypergeometric distribution analysis (p.adj.) is given.

| GO.ID | Description | DE | Total | p.adj. |
|---|---|---|---|---|
| GO:0055114 | Oxidation-reduction process | 15 | 490 | 0.001 |
| GO:0005506 | Iron ion binding | 8 | 100 | 7.94E-05 |
| GO:0020037 | Heme binding | 7 | 139 | 0.007 |
| GO:0016705 | Oxidoreductase activity, acting on paired donors, with incorporation or reduction of molecular oxygen | 6 | 62 | 0.001 |
| GO:0004497 | Monooxygenase activity | 4 | 45 | 0.023 |

**Table S8.** Significantly depleted biochemical pathways identified by their gene ontology (GO) terms in F0-exposed animals compared to controls. The number of significantly up-regulated (DE) and the total number of genes (Total) with this GO-term assigned, as well as the Bonferroni-adjusted p-value for the hypergeometric distribution analysis (p.adj.) is given.

| **GO.ID** | **Description** | **DE** | **Total** | **p.adj.** |
|---|---|---|---|---|
| GO:0005762 | mitochondrial large ribosomal subunit | 5 | 31 | 0.003 |
| GO:0071011 | precatalytic spliceosome | 4 | 16 | 0.003 |
| GO:0071013 | catalytic step 2 spliceosome | 5 | 34 | 0.004 |
| GO:0070125 | mitochondrial translational elongation | 2 | 2 | 0.013 |
| GO:0003723 | RNA binding | 12 | 295 | 0.020 |
| GO:0005686 | U2 snRNP | 3 | 14 | 0.050 |

### Differentially expressed and epigenetically relevant genes

**Table S9.** Differentially expressed genes in F0-exposed animals compared to controls that code for epigenetically relevant genes. GeneID, log2 fold change and adjusted p-value for the statistical comparison as well as a description and reference for the relation of the gene to epigenetic mechanisms is given.

| **GeneID** | **Log2FC** | **p-adj** | **Description** | **references** |
|---|---|---|---|---|
| DMV1G035110T0 | -13 | 0.047 | methyltransferase 6 | affect the epigetic landscape by histone methylation (Guccione et al., 2007) |
| DMV1G054830T0 | 23 | 0.017 | JmjC domain-containing histone demethylation protein | Tsukada et al. ( 2006) |
| DMV1G070960T0 | -22 | 0.040 | histone deacetylase 8 | Van Den Wyngaert et al. (2000) |
| DMV1G075340T0 | -13 | 0.037 | Chromatin-remodeling complex ATPase chain Iswi | Bartholomew (2014) |

**Data File S1 (separate file).** Differential gene expression analysis for pairwise comparisons between F0-exposed vs. control, Exposed vs. Control and Exposed vs F0-exposed. Tables list the gene ID based on the reference genome, a gene description, log 2 fold change for the comparison, standard error of the fold change (lfcSE), adjusted p-value (padj), the sequence length, the e-value and finally the GO IDs and GO-names
